# Cbp1-Cren7 chromatinization of CRISPR arrays favours transcription from leader-over cryptic promoters

**DOI:** 10.1101/2023.03.24.534125

**Authors:** Fabian Blombach, Michal Sýkora, Jo Case, Xu Feng, Diana P Baquero, Thomas Fouqueau, Duy Khanh Phung, Declan Barker, Mart Krupovic, Qunxin She, Finn Werner

## Abstract

CRISPR arrays form the physical memory of CRISPR adaptive immune systems by incorporating foreign DNA as spacers that are often AT-rich and derived from viruses. As promoter elements such as the TATA-box are AT-rich, CRISPR arrays are prone to harbouring cryptic promoters. Sulfolobales harbor extremely long CRISPR arrays spanning several kilobases, a feature that is accompanied by the CRISPR-specific transcription factor Cbp1. Aberrant Cbp1 expression modulates CRISPR array transcription, but the molecular mechanisms underlying this regulation are unknown. Here, we characterise the genome-wide Cbp1 binding at nucleotide resolution and characterise the binding motifs on distinct CRISPR arrays, as well as on unexpected non-canonical binding sites associated with transposasons. Cbp1 recruits Cren7 forming together ‘chimeric’ chromatin-like structures at CRISPR arrays. We dissect Cbp1 function *in vitro* and *in vivo* and show that the third HTH domain is responsible for Cren7 recruitment, and that Cbp1-Cren7 chromatinization plays a dual role in the transcription of CRISPR arrays. It suppresses spurious transcription from cryptic promoters within CRISPR arrays but enhances CRISPR RNA transcription directed from their cognate promoters in their leader region. Our results show that Cbp1-Cren7 chromatinization drives the productive expression of long CRISPR arrays.

## Introduction

Chromatinization of DNA regulates transcription in all domains of life, including eukaryotes, bacteria and archaea. Chromatin proteins can regulate transcription by modulating the access of transcription factors and RNA polymerase (RNAP) to the gene promoter, but also suppress transcription from cryptic promoters inside genes which in particular in the antisense direction can be detrimental to productive gene expression.

In *Salmonella* and *Escherichia coli*, the chromatin protein H-NS preferentially binds to AT-rich DNA that is enriched in cryptic promoter elements including the Pribnow box, or ‘−10’ element that is AT-rich [1]. Deletion of H-NS in *E. coli* leads to recruitment of RNA polymerase to these cryptic promoters thereby depleting the pool of free RNA polymerase [2].

CRISPR-Cas systems evolved as adaptive immune system of prokaryotes against mobile genetic elements. The physical memory of this system is formed by CRISPR arrays, genomic regions that encompass clusters of repeat sequences interspersed by spacers derived from foreign DNA [3]. Transcription of these CRISPR arrays produces pre-crRNAs that are further processed into crRNA units encompassing single spacer-repeat units. Pre-crRNAs are amongst the longest non-coding RNAs in prokaryotes next to the 16S and 23S rRNAs. In *E. coli*, dedicated antitermination complexes ensure processivity of CRISPR array transcription by preventing premature Rho-dependent transcription termination similar to the antitermination complexes facilitating rRNA transcription [4]. Beyond *E. coli*, little is known about specific mechanisms controlling CRISPR array transcription. Sulfolobales harbour a number of general chromatin proteins, in particular Alba, Sul7 and Cren7 [5]. Initial chromatin immunoprecipitation sequencing (ChIP-seq) data suggested increased occupancy of Cren7 at CRISPR arrays in *Saccharolobus solfataricus* but whether this affects CRISPR function remains unknown [6].

Members of the Sulfolobales order have multiple CRISPR systems and multiple CRISPR arrays spanning often > 100 spacers and resulting in pre-crRNAs of several kilobases in length. The extraordinary length of CRISPR arrays in Sulfolobales coincides with the presence of the <u>C</u>RISPR array <u>b</u>inding <u>p</u>rotein <u>1</u>, Cbp1, that binds to the repeat sequences in CRISPR arrays [7, 8]. Cbp1 comprises three helix-turn-helix (HTH) domains that are derived from domain duplications [7, 8].

Deletion of *cbp1* in *Sulfolobus islandicus* leads to reduced levels of pre-crRNA and processing intermediates, while Cbp1 overexpression increases the levels of pre-crRNA [7]. These observations lead to the concept that Cbp1 is a positive transcription elongation factor specific for CRISPR arrays [7], which is highly counterintuitive as a protein binding to numerous sites downstream of an elongating RNAP likely acts as a ‘roadblock’ factor. An alternative, apparently mutually exclusive hypothesis suggests that the regulated binding of Cbp1 to CRISPR arrays might induce premature transcription termination to adjust crRNA levels in a vectorial fashion, i. e. favouring the synthesis of crRNAs in the upstream segments of the arrays [9]. New spacers that provide greater protection against the current viral pool are generally integrated at the 5’ end of CRISPR arrays. Premature transcription termination enriches the crRNAs bearing new spacers in the total crRNA pool and counteract the ‘dilution effect’ arising from the transcription of distal spacers in very long CRISPR arrays.

Previous data suggested that Cbp1 suppresses spurious transcription from internal promoters [7]. Cbp1 could also have additional functions, e. g. it could facilitate the regulation of CRISPR systems in response to infection, e. g. during spacer acquisition. Infection with the SIRV2 [10] and STSV2 [11] viruses induces upregulation of CRISPR array expression in *Sulfolobus islandicus*, but it remains unknown whether this coincides with altered Cbp1 chromatinization of CRISPR arrays.

Here we provide a genome-wide yet detailed characterisation of Cbp1 function in CRISPR array transcription that enhances our understanding of the regulation of CRISPR systems and provides more general insights into the role of unorthodox chromatin-like proteins in transcription regulation in prokaryotes. We have mapped Cbp1 binding patterns globally and at nucleotide resolution, dissected the binding motifs and modes on distinct arrays, and tested the impact of Cbp1 binding on transcription *in vitro* and *in vivo*. We show that Cbp1 directly recruits Cren7 to the 3’-end of CRISPR repeats suppressing transcription initiation from cryptic promoters in spacers while facilitating transcription from CRISPR leader promoters. Additional binding sites of Cbp1 associated with transposases and the leaders of alternative CRISPR arrays hint on a wider regulatory function of Cbp1 linking defense systems and mobile genetic elements.

## Results

### Cbp1 recruits Cren7 to chromatinize CRISPR arrays

To gain insight into the chromatinization of CRISPR arrays *in vivo* and to test whether Cbp1 and Cren7 chromatinization of CRISPR arrays are interdependent, we determined the genome-wide occupancy of Cbp1 and Cren7 in *S. solfataricus* strain P2 by ChIP-seq. The *S. solfataricus* genome harbours six CRISPR arrays labelled A to F. All six CRISPR arrays showed increased ChIP-seq occupancy for both Cbp1 and Cren7 (Figure 1a). The six CRISPR arrays can be classified into three groups based on their CRISPR repeat sequences with A/B and C/D having identical repeats, respectively, while E/F repeats differ by a single nucleotide (Figure 1b). Notably, Cbp1 and Cren7 occupancy was correlated between the groups of CRISPR arrays (noticeable as distinct clusters Figure 1c) as well as within each cluster formed by each CRISPR array group (Figure 1c).

**Figure 1:**
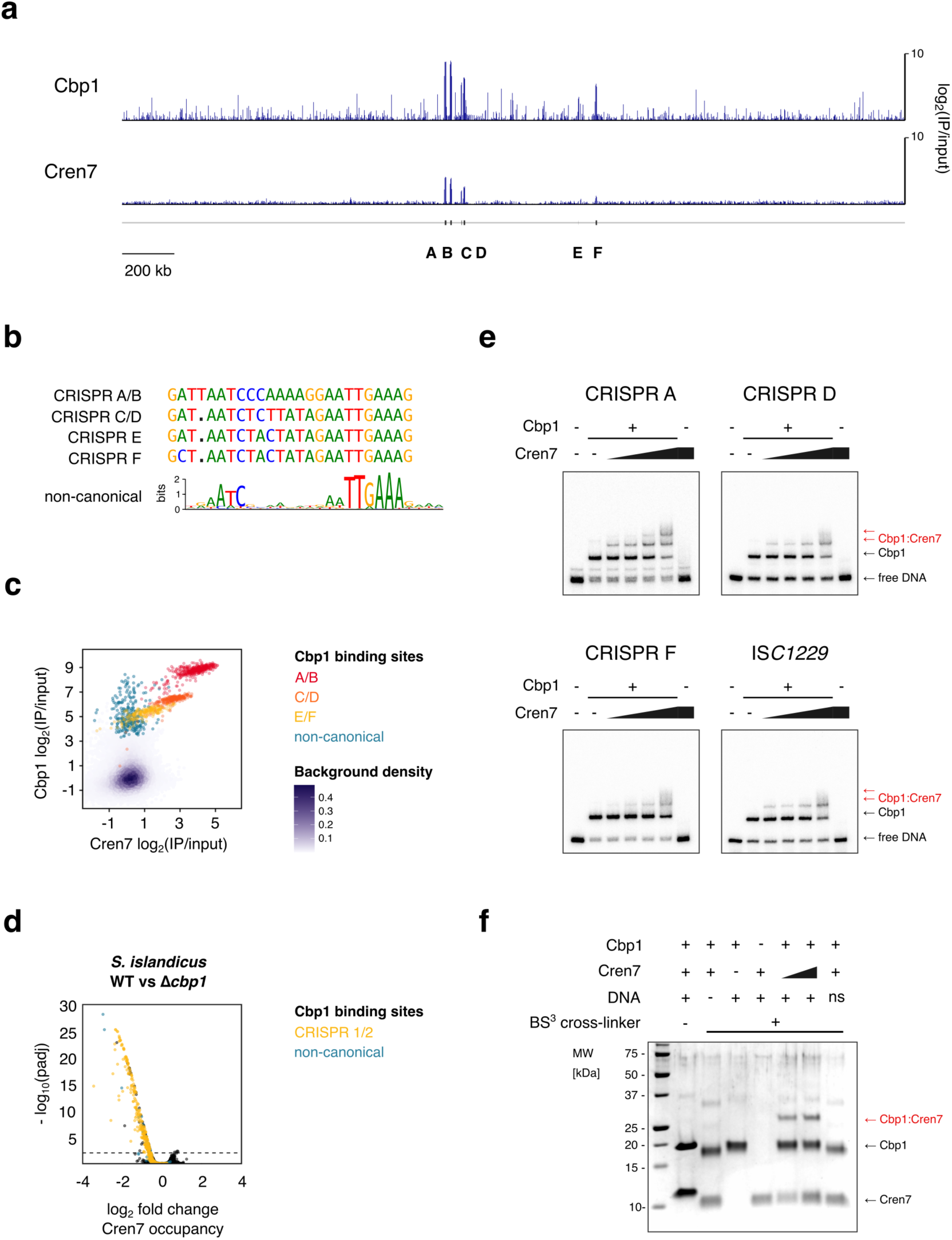
Cbp1 recruits Cren7 to chromatinise CRISPR arrays. (a) Genome-wide overview of Cbp1 and Cren7 occupancy on the *S. solfataricus* P2 genome. The location of the six CRISPR arrays is indicated. ChIP-seq data represent the mean of two biological replicates with 2000 bp bin size. (b) Sequence alignment of the different repeat sequences in the *S. solfataricus* P2 genome and WebLogo of the motif identified in non-canonical Cbp1 binding sites(n=92). (c) Scatter plot depicting correlated occupancy of Cbp1 and Cren7 on CRISPR arrays. Average occupancy (normalised against input) was calculated for 30 bp consecutive bins for two biological replicates. (d) Cren7 recruitment to CRISPR arrays is reduced in a *cbp1* deletion strain. Volcano plot showing the differential binding analysis of Cren7 ChIP-seq data for *S. solfataricus* E234 (WT) and Δ*cbp1*. Read counts were calculated for 30 bp consecutive bins for two biological replicates. (e) Cbp1 facilitates recruitment of Cren7. EMSA testing Cbp1 and Cren7 binding to three different CRISPR repeats and a non-canoncial binding site of Cbp1. 12.5 nM Cbp1 and 3.125 to 25 nM Cren7 (2x dilution series) were incubated with 5’-radiolabelled dsDNA template encompassing the CRISPR repeats and 20 bp flanking spacer DNA on either side or the corresponding region for a non-canonical binding site derived from a IS*C1229* transposon. Representative gels of three technical replicates are shown. (f) Cbp1 and Cren7 directly interact with each other in a DNA-dependent manner. Cbp1:Cren7:CRISPR A repeat DNA complexes were assembled with 2.5 µM Cbp1, 2.5 or 5 µM Cren7, and 2.5 µM dsDNA template. The complexes were then subjected to protein:protein crosslinking with 1 mM BS^3^. Crosslinked samples were resolved by SDS-PAGE with Coomassie staining. A Cbp1:Cren7 crosslinked species of ~25 kDa (labelled in red) was formed strictly in the presence of BS^3^, Cbp1, Cren7 and the CRISPR A repeat 1 template. Replacement of the CRISPR DNA template with a non-specific control DNA did not yield any detectable Cbp1:Cren7 crosslinked species. Some background signal at ~37 kDa can be attributed to a small fraction of Cbp1 dimers.

In addition to the CRISPR array associated binding of Cbp1, we also identified several non-canonical binding sites for Cbp1 (92 peaks with at least five-fold enrichment of Cbp1). These binding sites featured a 21 bp motif with a consensus sequence matching the CRISPR repeat consensus sequence with the strongest sequence conservation at six base pairs close to the 3’-end of the repeats (Figure 1b). Notably, these non-canonical Cbp1 binding sites in *S. solfataricus* did generally not show any enrichment of Cren7 compared to the genomic background (Figure 1c).

To test whether there is a causal relationship between Cbp1 and Cren7 chromatinization of CRISPR arrays, we tested Cren7 occupancy on CRISPR arrays in a *cbp1* deletion strain generated in *S. islandicus* REY15A strain E234, a genetically tractable strain closely related to *S. solfataricus* [7]. Similar to our *S. solfataricus* results, both Cbp1 and Cren7 chromatinized the two CRISPR arrays in E233S. Even though Cren7 expression levels are not affected by *cbp1* deletion (Supplementary Figure 1b), the Cren7 occupancy on the two CRISPR arrays was severely reduced (Figure 1d). Deletion of Cbp1 is accompanied by a deletion of a ~28 kb genomic region between two IS*200*/IS*605* family transposases, SiRe_0633 (SIRE_RS03230) and SiRe0665 (SRE_RS03390). However, this deletion is unlikely to affect Cren7 binding (see Supplementary table 1).

In summary, our ChIP-seq data thus suggest that Cbp1 enhances or facilitates Cren7 recruitment. We tested this hypothesis directly in electrophoretic mobility shift assays (EMSA) on dsDNA templates with a single CRISPR repeat from CRISPR arrays A, D and F or a non-canonical binding site derived from an IS*C1229* transposon. The EMSAs confirmed Cbp1-dependent recruitment of Cren7 to all DNA templates. Notably, Cbp1-bound CRISPR A repeats appeared to have an overall higher binding affinity for Cren7 and at higher Cren7 concentrations a second Cren7 monomer was recruited to Cbp1-bound CRISPR A repeats (Figure 1e). Cbp1 itself appeared to show weaker affinity for the CRISPR A repeat than for the CRISPR F repeat in direct contrast to our ChIP-seq occupancy data but in line with previous findings [7]. The Cren7 concentrations used in our experiments were below the range where Cren7 efficiently chromatinizes DNA [12] explaining the absence of detectable DNA-binding by Cren7 in the absence of Cbp1.

Recruitment of Cren7 could be mediated by direct physical interaction between Cbp1 and Cren7 or by Cbp1-induced topological changes in the DNA template facilitating Cren7 binding. To test whether there is a direct physical interaction, we conducted crosslinking assays using the amine-specific crosslinker Bis(sulfosuccinimidyl)suberate (BS^3^). We observed a crosslinked species corresponding to the expected combined molecular weight of Cbp1 and Cren7 dependent on the presence of Cbp1, Cren7, and a DNA template bearing a CRISPR repeat (Figure 1f). The crosslinked species was not observed in control experiments with a non-specific DNA template (Figure 1f). Our data thus demonstrate that Cbp1 recruits Cren7 through direct physical interactions.

### The essential nature of Cren7 does not depend on Cbp1

C*ren7* is essential for *S. islandicus* viability but Cbp1 is not [7, 13]. Deletion of *cbp1* coincides with loss of Cren7 binding to its main target in the genome, the CRISPR arrays. To test a genetic interaction between the two chromatin proteins, we attempted to delete *cren7* (SiRe_1111) in the Δ*cbp1* strain. However, we were unable to delete *cren7* in *S. islandicus* Δ*cbp1* whereas the non-essential gene SiRe_0782 [13] that serves as positive control could be deleted (Supplementary Figure 1c). We conclude that the lethality of the *cren7* deletion cannot be compensated for by concomitant deletion of *cbp1*, which indicates that the essentiality of Cren7 is not connected to CRISPR array chromatinization.

### Architecture of the Cbp1-Cren7-DNA complex

To gain insight into the topology of the Cbp1-Cren7-DNA complex, we used ChIP-exo, a method combining ChIP-seq with 5’→3’ exonuclease trimming, to map the footprints of Cbp1 and Cren7 on CRISPR repeats at nucleotide resolution. Aggregate profiles for Cbp1 footprints on CRISPR repeats of arrays A and B revealed a 5’ border at position −5 relative to the start of the CRISPR repeat and a 3’ border located at position +16 downstream of the repeat start (Figure 2a). Notably, position +16 is upstream of the region encompassing the core Cbp1 binding motif (Figure 1b). Cbp1 footprints for the non-canonical Cbp1 binding sites showed a similar pattern with the 3’ border located at the corresponding position immediately upstream of the 3’-terminal core binding motif (Supplementary Figure S2). Because of this lack of protection over the core binding motif, the Cbp1 ChIP-exo footprints appear not to represent the full extent of the Cbp1 binding site and are potentially biased by the cross-linkability at different positions. The corresponding ChIP-exo profiles for Cren7 were highly similar to those obtained for Cbp1 potentially due to stronger protein-protein crosslinking relative to DNA-protein crosslinking at our experimental conditions consistent with other ChIP-exo data we obtained previously [14]. Nevertheless, the Cren7 aggregate profile showed additional protection downstream of position +16 downstream reaching into the 3’-flanking spacer sequence. Combined with the pattern of sequence conservation (Figure 1b), the ChIP-exo data suggest that Cbp1:Cren7 binding covers the entire CRISPR repeat and stretches downstream into the flanking spacer.

**Figure 2:**
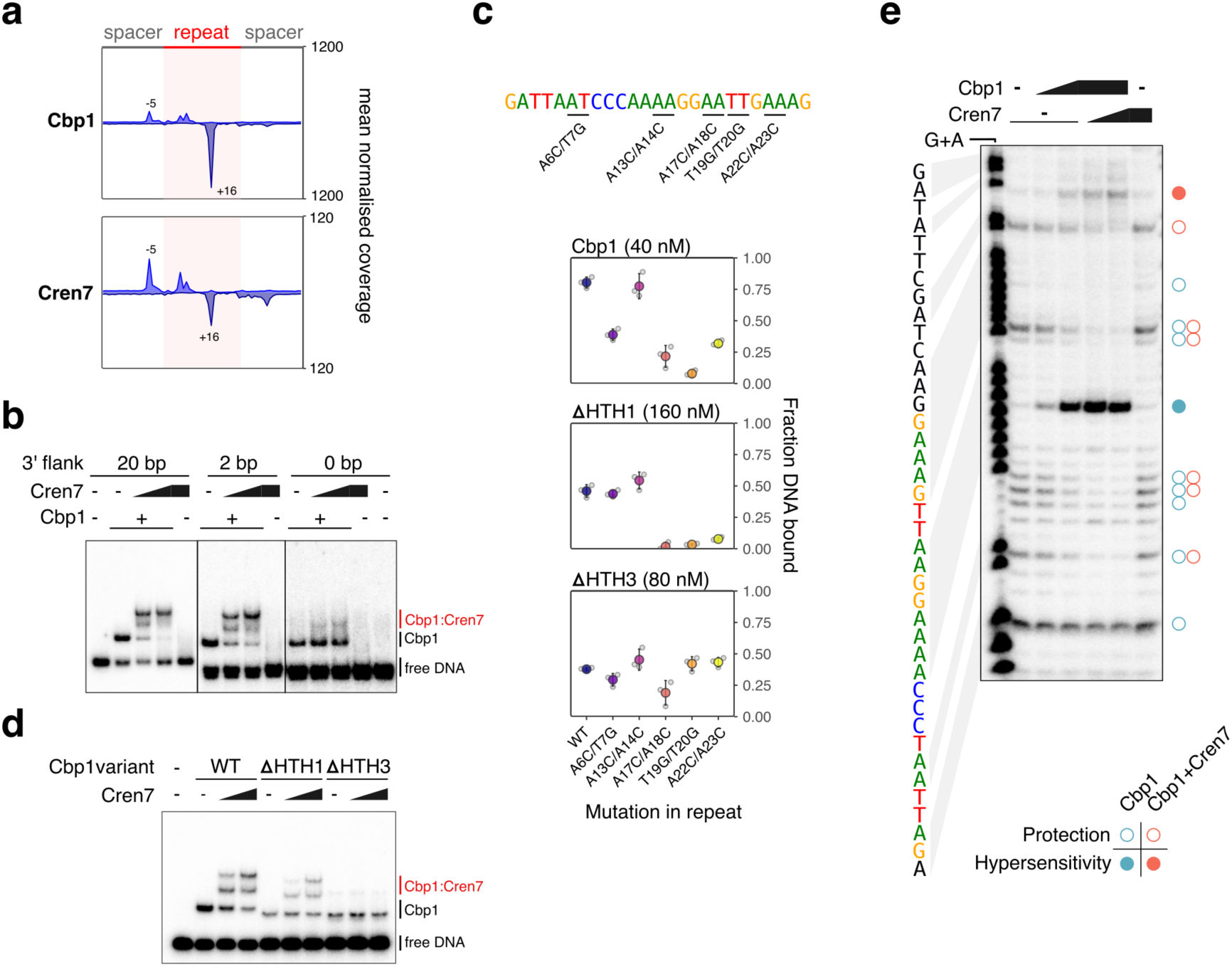
Architecture of Cbp1:Cren7 chromatinization. (a) Aggregate plots of Cbp1 and Cren7 ChIP-exo occupancy over 208 repeats from CRISPR arrays A and B. Signal for the plus and minus strand (relative to CRISPR array orientation) is plotted above and below the x-axis, respectively. The aggregate mean signal was calculated from the geometric mean of two biological replicates scaled to reads per million. (b) Cren7 recruitment to Cbp1:CRISPR DNA complexes depends on the flanking DNA downstream of the CRISPR repeat. EMSAs with templates bearing a CRISPR A repeat with 20 bp upstream and 20, 2 or 0 bp downstream flanking sequence. The free DNA of the shorter templates appeared as double-band, possibly due to some denaturation at the incubation temperature of 75 °C. (c) Orientation of Cbp1 binding to CRISPR repeats. Binding of Cbp1 and HTH-deletion mutants ΔHTH1 and ΔHTH3 to mutated CRISPR repeat sequences was assessed by EMSAs and the fraction of DNA bound by Cbp1 was plotted (mean of three biological replicates and standard deviation are shown). Double mutations were introduced into the CRISPR repeat sequence based on regions showing stronger sequence bias in the motif identified for non-canonical Cbp1 binding sites (Figure 1B). The A13C/T14G double mutant was included as control for a region of CRISPR repeats that shows no sequence bias in non-canonical binding sites. To compensate for the overall lower affinity of the ΔHTH1 and ΔHTH3 variants, their concentration was raised to 160 and 80 nM, respectively. (d) Cren7 recruitment to Cbp1:CRISPR repeat complexes depends on the HTH3 domain of Cbp1. EMSA assay testing Cren7 recruitment to CRISPR DNA-bound Cbp1 (12.5 nM), ΔHTH1 (50 nM) and ΔHTH3 (25 nM). We observed a low level of recruitment of a second Cbp1 ΔHTH3 that was also present at higher concentrations of WT Cbp1 (data not shown). (e) DNase foot-printing assays corroborate Cbp1-dependent Cren7 deposition at the downstream spacer. Foot-printing assays were carried out with 12.5 or 25 nM Cbp1 and 25 or 50 nM Cren7. A G+A sequencing ladder is shown on the left as reference. Cbp1 and Cbp1:Cren7-induced protection against DNase cleaveage is indicated with open circles and hypersensitivity is indicated with full circles.

To test the role of the downstream DNA flanking the core motif in Cbp1 and Cren7 binding *in vitro*, we conducted EMSAs with DNA templates including varying lengths of 3’-flanking spacer sequences. We observed recruitment of two Cren7 monomers was retained when the 3’-flanking DNA was trimmed from 20 bp to 2 bp (Figure 2b). Further trimming of the remaining flanking DNA abolished Cren7 recruitment in line with Cren7 interacting with the 3’ flanking DNA (Figure 2b).

### Dissection of Cbp1 HTH domain contributions to DNA binding

Cbp1 is composed of three HTH domains connected by short linker regions, these domains are homologous as the protein likely arose by a domain duplication mechanism. To determine how the HTH domains contribute to CRISPR repeat binding, we deleted either the first or the third HTH domain (ΔHTH1 and ΔHTH3). Both deletion variants were expressed at comparable levels to the wild type full length protein, and their thermostability validated that the mutations had not impaired correct protein folding (data not shown). We tested the relative orientation of the Cbp1 HTH domains on the CRISPR repeats by combining mutations in the CRISPR repeat with the Cbp1 ΔHTH1 and ΔHTH3 variants. If the N- or C-terminal HTH domain interacts with a specific region in the CRISPR repeat, we reasoned that mutations in this region should not affect binding of a Cbp1 variant where the HTH is absent. We tested four different double transversion mutations in the CRISPR repeat at positions corresponding to positions with strong sequence conservation in the Cbp1 binding motif: A6C/T7G, A17C/A18C, T19G/T20G, A22C/A23C. As a control, we also included the A13C/T14G mutation, a region without any apparent sequence conservation in the core motif. We first tested the effect of these mutations on full-length Cbp1. All mutations but the A13C/T14G control reduced Cbp1 binding relative to the WT repeat sequence (Figure 2c). Both Cbp1ΔHTH1 and Cbp1ΔHTH3 showed overall weaker binding. Crucially, Cbp1ΔHTH1 appeared to be unaffected by the 5’-terminal A6C/T7G mutation whereas the 3’-terminal mutations drastically reduced binding. Conversely, Cbp1ΔHTH3 binding appeared to be unaffected by the 3’-terminal two mutations (Figure 2c). Our data suggest that the three HTH domains of Cbp1 align along the CRISPR repeat in 5’ to 3’ direction. We corroborated these results by testing Cren7 recruitment by the HTH-deletion variants in EMSAs. In line with HTH3 binding to the 3’-terminal core motif where Cren7 is recruited, the Cbp1ΔHTH3 mutant failed to recruit Cren7 (Figure 2d).

We corroborated our data by carrying out DNase I footprinting experiments with Cbp1 and Cren7. Previous DNase I footprinting experiments for Cbp1 binding to CRIPSR repeats revealed a DNase hypersensitivity site in the centre of the repeat flanked by protected regions [8]. Our data reproduced these findings (Figure 2e). The addition of Cren7 enhanced the protection and extended it at least 6 bp into the spacer downstream of the repeat. A new minor hypersensitivity site appeared at position 10 in the downstream spacer(Figure 2e). Taken together, our data establish the overall topology of the Cbp1-Cren7-DNA complex with Cren7 being recruited to the 3’-terminal region of the CRISPR repeat covering in part the flanking spacer.

### Cbp1-Cren7 chromatinization of CRISPR arrays suppresses spurious transcription from cryptic promoters

Having established the topology of Cbp1-Cren7-DNA complex, we set out to investigate how Cbp1-Cren7 chromatinization affects transcription of CRISPR arrays. We compared transcription of *S. solfataricus* P2 CRISPRB and CRISPRF that show high and low levels of Cbp1-Cren7 chromatinization, respectively. To probe how Cbp1-Cren7 chromatinization relates to the recruitment of the transcription machinery, we mapped the genome-scale occupancy of transcription initiation and elongation factors, as well as RNAP and regulatory factors using our previously published ChIP-seq data [14]. To complement these binding data with transcription output, we analysed short RNA sequencing data generated by Cappable-seq [14], a method that is highly selective for 5’-triphosphorylated RNA and thus capable to detect transcription initiation events with high sensitivity [15].

Our data revealed several TSSs associated with internal promoters both in sense and antisense orientation within CRISPR array F (Figure 3a). Basal transcription factors including TBP, TFB and TFE form transcription preinitiation complexes with RNAP and facilitate promoter-directed transcription initiation. During early elongation, proximal to the promoter, the elongation factors Spt4/5 and Elf1, and the termination factor aCPSF1, are recruited to the transcription elongation complex [14, 16]. The heterogenous factor composition and occupancy detected at the internal promoters reflect the RNAP in different stages of the transcription cycle. E. g. the promoters in array B showed ChIP-seq occupancy of TFB and RNAP, but not Spt4/5, suggesting that these promoters allow only for preinitiation complex formation (Figure 3a). In contrast, several promoters within array F showed occupancy of TFB, RNAP, Spt4/5 and aCPSF1 suggesting that RNAP had progressed to the transcription elongation complex and possibly even premature transcription termination. The ChIP-seq occupancy corresponded overall well with the detection of 5’-triphosphorylated RNAs by Cappable-seq at these promoters. The internal cryptic promoters generally coincided with local minima in Cren7 ChIP-seq occupancy. These data suggest that Cbp1-Cren7 chromatinization might compete with transcription from cryptic promoters within CRISPR arrays.

**Figure 3:**
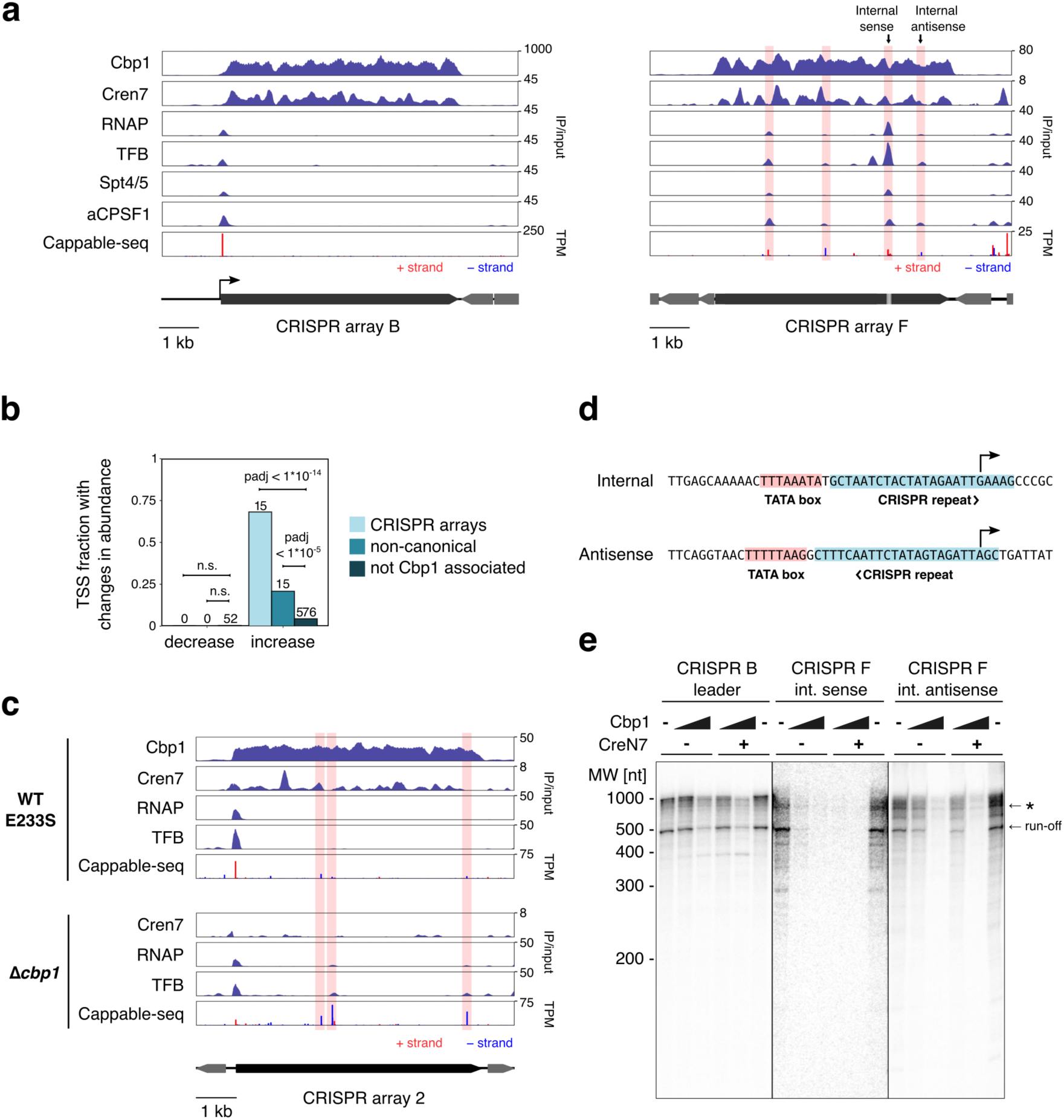
Cbp1:Cren7 chromatinization of CRISPR arrays prevents spurious transcription from internal promoters. (a) *S. solfataricus* CRISPR arrays B and F with high and low Cbp1:Cren7 occupancy, respectively, show different levels of spurious transcription. ChIP-seq occupancy profiles for Cbp1, Cren7, RNAP, initiation factor TFB, elongation factor Spt4/5 and termination factor aCPSF1 are shown. The mean of two biological replicates is shown with the range depicted as semi-transparent ribbon. The lower panel shows Cappable-seq data as 5’-end coverage for 5’-triphosphorylated short RNAs (~20-200 nt). The geometric mean from two biological replicates was scaled to transcripts per million (TPM). The position of the F1 internal and antisense promoters tested in *in vitro* transcription (panel D) are indicated. (b) Deletion of *cbp1* leads to activation of TSSs within CRISPR arrays and close to non-canoncial binding sites of Cbp1. TSSs were mapped using Cappable-seq for *S. islandicus* REY15A and *cbp1* deletion strains 2 and 3. Differential expression of TSSs (padj < 0.01) was identified using Deseq2. Enrichment of differentially expressed TSSs close to Cbp1 binding sites was assessed using Fisher’s exact test with Bonferroni multiple testing correction. (c) *cbp1* deletion elevates spurious transcription from *S. islandicus* REY15A CRISPR arrays 1 and 2. ChIP-seq occupancy profiles for Cbp1, Cren7, RNAP, and TFB alongside Cappable-seq data for the 5’-end coverage of triphosphorylated short RNAs are shown for parental strain E234 and the *cbp1* deletion strain [7]. (d) Sequence of the internal sense and antisense promoters in CRISPR F1 (see panel a), respectively used in cell-free transcription experiments. (e) Cbp1 and Cbp1:Cren7 chromatinization prevents spurious transcription *in vitro.* The internal sense and antisense promoters in CRISPR F1 (see panel A) were tested in cell-free transcription assay with increasing concentrations of recombinant Cbp1 added (0, 100, 300 nM) and in presence (300 nM) or absence of recombinant Cren7. Affinity-purified radiolabelled transcripts were resolved on a denaturing polyacrylamide gel. A representative gel of three technical replicates is shown. The position of the run-off transcripts (516 nt for CRISPR B, 505 nt for CRISPR F promoters) is indicated. The asterisk denotes longer, unspecific transcripts appearing with longer incubation times.

To establish a causal relationship between Cbp1 chromatinization and the suppression of spurious transcription, we compared transcription profiles for the *S. islandicus Δcbp1* strain with the parental strain E233S by Cappable-seq RNA sequencing as well as ChIP-seq for RNAP and TFB. As in *S. solfataricus*, cryptic promoters appeared to be active in spacers with locally decreased Cren7 occupancy in E233S. To map changes in transcription activity of these cryptic promoters, we conducted a differential expression analysis of the Cappable-seq data for the identified TSSs in the *S. islandicus cbp1* deletion and the parental strain. To avoid the additional deletion of a genomic region in our original Δ*cbp1* strain, we reconstructed the strain. The two biological replicates for the parental E233S strain as well as two independent Δ*cbp1* strains correlated well across the quantified signal for 13150 TSSs (Spearman’s r=0.97 for the E233S replicates and r=0.96 for the two Δ*cbp1* strains, Supplementary Figure 3a). The differential expression analysis confirmed our hypothesis as the *cbp1* deletion results in substantial changes in TSS utilisation and transcriptome changes with 658 out of 13,150 TSSs differentially transcribed (padj < 0.01, Figure 3b). The TSSs associated with leader promoters that direct CRISPR array transcription appeared to be downregulated by the deletion of *cbp1* (log_2_-fold change of −1.56, padj <0.053 for CRISPR1 and −1.7, padj < 1*10^−15^ for CRISPR2) and in good agreement with the reduced level of pre-crRNA observed in Δ*cbp1* using Northern blots [7]. This effect could be caused by a reduced transcription initiation frequency or by a feedback mechanism from slow/paused transcription elongation complex interfering with transcription preinitiation complexes and productive initiation. Next, we investigated the effect of the Cbp1 deletion on cryptic promoters inside CRISPR arrays. A large fraction of TSSs and promoters residing within the two CRISPR arrays were transcribed at higher levels upon *cbp1* deletion (15 out of 22, padj < 1*10^−14^, multi-comparison Fisher’s exact test, Benjamini-Hochberg correction), in good agreement with the hypothesis that Cbp1 binding suppresses internal and spacer promoter transcription. Non-canonical binding sites of Cbp1 that are not associated with CRISPR arrays featured a repeat-like binding motif consistent with *S. solfataricus* (Supplementary Figure 3b). TSSs associated with non-canonical Cbp1 binding sites, i. e. within 50 bp of Cbp1 peak summit, also showed a slightly higher fraction with increased expression after *cbp1* deletion (15 out of 72, padj < 1*10^−5^). ChIP-seq data for RNA polymerase and TFB in the E233S WT and the Δ*cbp1* strain demonstrate that the increased transcription resulted from cryptic promoters active already in the WT strain. Thus, spurious transcription from CRISPR-array internal promoters appears to be suppressed by Cbp1:Cren7 chromatinization (Figure 3c).

We previously described that global mRNA levels in archaea are correlated with the fraction of RNAPs that successfully ‘escape’ into the productive elongation phase of transcription. During normal growth, RNAP is predominantly located in the promoter-proximal regions of CRISPR arrays (Figure 3a) [14]; the release from this state, an increase of RNAP escape, likely triggers up-regulation of CRISPR expression in response to viral infection [10, 11]. Notably, promoter-proximal peaks of RNA polymerase were also observed in the Δ*cbp1* strain indicating that RNA polymerase accumulation is not strongly dependent on Cbp1 chromatinization (Figure 3c).

To test the competition between Cbp1 binding and transcription directly, we used cell-free *in vitro* transcription assays to compare two internal promoters within *S. solfataricus* CRISPR array F (Figure 3a), one in sense and one in antisense orientation, with the leader promoter of CRISPR array B that directs pre-crRNA transcription. The internal sense and antisense promoters are both overlapping with CRISPR repeats that are downstream of their TATA-boxes and cover the TSSs (Figure 3d). The DNA templates used in the cell-free transcription assays encompassed a region spanning −100 to +500 relative to the TSS (+511 for CRISPR B) plus five GC base pairs at either end.

The levels of endogenous Cbp1 in the cell lysate are insufficient to saturate the large excess of DNA template added to the reaction, allowing us to manipulate the Cbp1-chromatinization of the templates by adding recombinant Cbp1 (Supplementary Figure 3c) while having the full complement of transcription initiation and elongation factors and RNA polymerase present in the lysate. All three promoters directed the synthesis of run-off transcripts of the expected size (Figure 3e), while no run-off transcript was detected in negative control reactions including TATA-box mutations of either promoter, confirming that the observed transcripts were promoter-specific (Supplementary Figure 3d). The addition of Cbp1 inhibited transcription of internal sense and antisense promoters 100 nM Cbp1 with total with near-total inhibition at 300 nM Cbp1. In contrast, the leader promoter remained largely unaffected at 100 nM Cbp1 with some repression at 300 nM Cbp1 (Figure 3e). Addition of Cren7 did not compound the repressive effect of Cbp1.

### Cbp1 enhances CRISPR array transcription in a minimal system

The experiments with the Cbp1 deletion strain suggest that Cbp1 stimulates CRISPR array expression. To test whether this effect is independent of the CRISRP leader promoters, we cloned chimeric transcription templates by fusing the well characterised SSV1 T6 model promoter to the first 511 bp of *S. solfataricus* CRISPR A and B arrays containing eight repeat sequences. To test a potential orientation bias of Cbp1, we compared templates with the CRISPR B fragment cloned in either orientation. EMSA experiments validated Cbp1 binding to these templates with a saturation of Cbp1 binding occurring at 300 nM (Supplementary Figure 4). To test Cbp1 stimulation under rigorously defined conditions, we carried out *in vitro* transcription assays in a minimal, reconstituted assay with recombinant transcription initiation factors. Here we ensure single-round transcription by synchronising initially transcribing complexes at register +6 by omitting CTP and UTP. RNA synthesis commences by the addition of CTP and UTP in the presence of a large excess of a TFB variant that inhibits RNAP reinitiation [14, 17, 18]. *In vitro* transcription using the CRISPR B templates resulted in the synthesis of 516nt run-off transcripts within 2 min with good processivity, i. e. only weak partial transcript patterns indicative of elongation pausing (Figure 4b). The addition of Cbp1 (300 nM) stimulated transcription, while addition of Cren7 had no effect on its own nor did it enhance the Cbp1 stimulation (data not shown). This stimulation could be observed using both templates, i. e. independently of the orientation of the CRISPR array and Cbp1-binding sites. The transcript profiles suggest that Cbp1 enhances the overall processivity of transcription. To rule out that direct contacts between the RNAP in the pre-initiation complex and the promoter-proximal Cbp1 increases transcription at the level of initiation, we tested a 5’ truncated template lacking the first repeat (Δrepeat1). Cbp1 stimulated transcription to a similar extent from both wild type and Δrepeat1 templates, arguing against a role of Cbp1 for transcription initiation. Crucially, a control transcription template encompassing a T6 promoter fusion to a ~500 bp region lacking Cbp1 binding sites did not show stimulation by Cbp1 (Figure 4d). Our data show that Cbp1 chromatinization enhances transcription processivity in a minimal transcription system in the absence of elongation factors.

**Figure 4:**
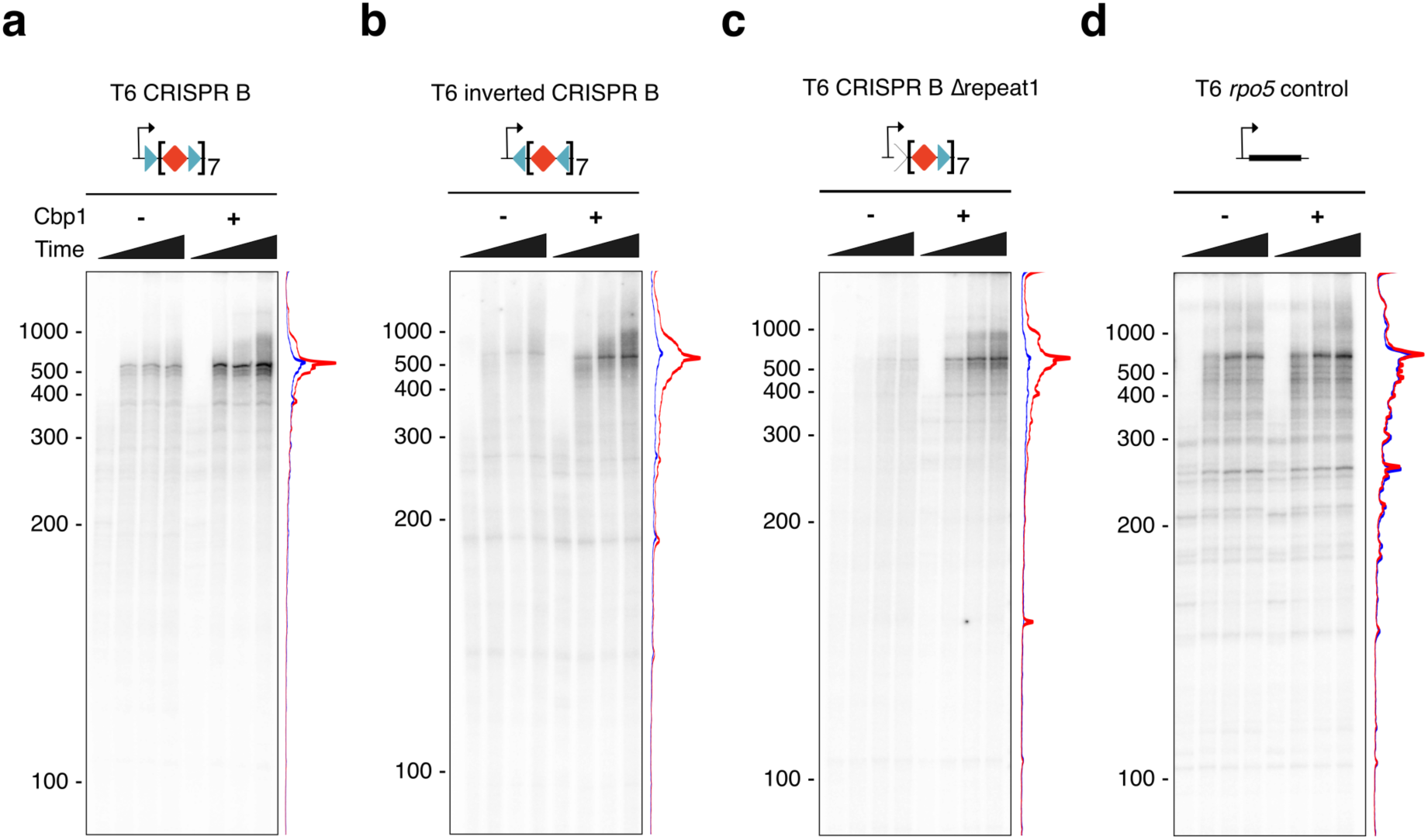
Cbp1 enhances CRISPR array transcription from leader promoters in a minimal transcription system. (a-d) Synchronised reconstituted *in vitro* transcription experiments with the strong T6 promoter fused to the first ~500 bp of *S. solfataricus* P2 CRISPR array B (a), the same CRISPR array sequence inverted (b), and the same sequence with the first CRISPR repeat randomised (c) in the presence or absence of 300 nM Cbp1. As control for a transcription template lacking Cbp1 binding sites we used a fusion of the T6 promoter to the *rpo5* gene from the archaeon *Methanocaldococcus janaschii* (d) [53]. Time points 1, 2, 3, and 4 min after release of RNAP from the promoter are shown. Lane profiles for time point 3 min with Cbp1 (red) or without (blue) are depicted on the right of each gel.

### Cbp1 and RNAP recruitment to CRISPR arrays in response to viruses

The expression of CRISPR arrays is induced by viral infection. This effect has been characterised by transcriptome analyses of *S. islandicus* LAL14/1 infected with the lytic virus SIRV2 [10]. *S. islandicus* LAL14/1 has five CRISPR arrays with two different, unrelated repeat sequences [19]. We used ChIP-seq of Cbp1 and RNA polymerase to (i) characterize Cbp1 binding to different *S. islandicus* CRISPR arrays, to (ii) probe for the presence of RNAP transcription complexes in spacers, and (iii) specifically test whether virus infection is accompanied by changes in Cbp1 and RNAP binding, by comparing uninfected with SIRV2 infected cells. CRISPR arrays 1 and 2 have repeat sequences identical to CRISPR F in *S. solfataricus* P2 (Figure 5a). In good agreement, our ChIP-seq results show that they are bound by Cbp1 (Figure 5b). In contrast, arrays 3, 4 and 5 have a divergent repeat sequence and are not bound by Cbp1 (Figure 5a,c). The CRISPR repeat sequences display differential prevalence in Sulfolobales species from different geographic locations, with the array 1/2-like repeat sequence being dominant in most analysed locations [20]. As expected, all arrays show a strong RNAP peak at the leader promoter corresponding to the PIC that facilitates their expression. However, while array 3, 4, and 5 show internal RNAP peaks indicative of RNAP recruitment to cryptic internal promoters, arrays 1 and 2 show no signs of any RNAP peaks within the array, similar to the CRISPR arrays in *S. islandicus* REY15A and *S. solfataricus* P2. These findings are consistent with Cbp1 chromatinization shielding CRISPR arrays against cryptic promoters. A detailed differential binding analysis of Cbp1 and RNAP between uninfected and infected cells was compromised by an increased genomic DNA background present in the ChIP-seq libraries after SIRV2 infection. We only observed subtle changes with the ChIP-seq occupancy of RNAP and Cbp1 within array 1, 3 and 4 estimated to be lower in the SIRV2-infected samples, probably as a result from higher genomic DNA background in the ChIP samples. However, SIRV2 infection induced expression of Cas operons (type I-A) previously discovered by RNA-seq [10] was well reflected in the ChIP-seq data with for example increased RNAP occupancy on a type I-A Cas operon (Figure 5b).

**Figure 5:**
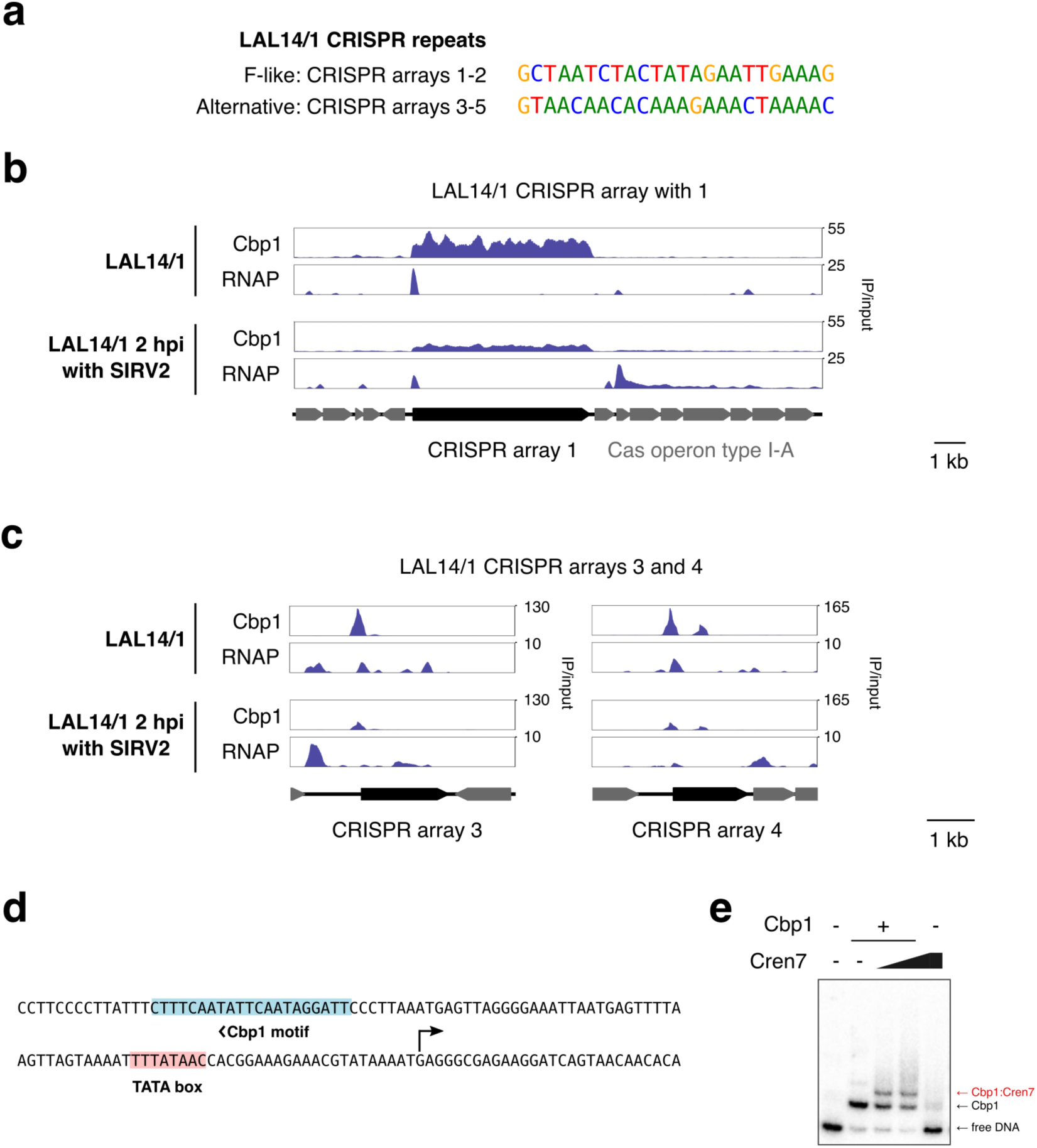
Cbp1 controls promoters of CRISPR arrays not chromatinized by Cbp1. (a) F-like and alternative CRISPR repeat sequences in *S. islandicus* LAL14/1 CRISPR arrays. (b) CRISPR arrays with F-like repeat sequences are Cbp1-chromatinized and chromatinization is maintained after SIRV2 infection. The mean signal of two biological replicates is shown. (c) Non-Cbp1-chromatinised CRISPR arrays are associated with Cbp1 peaks in their promoter region. (d) Sequence of the promoter of CRISPR array 4. A putative TSS and TATA-box were detected based on RNA-seq coverage of CRISPR transcripts [10]. (e) Cbp1 binds to the CRISPR 4 promoter *in vitro*. EMSA testing Cbp1 (6.25 nM) and Cren7 (12.5 or 25 nM) binding to a 5’-radiolabelled 128 nt dsDNA template bearing the CRISPR 4 leader promoter including a Cbp1 binding site.

Intriguingly, our ChIP-seq analysis identified a new class of strong Cbp1 binding sites in vivo, which were located upstream of the TATA/BRE motifs in the leader promoters in arrays 3, 4 and 5 (Figure 5d). Cbp1 and Cren7 bound to the upstream promoter sites in EMSAs (Figure 5e) but had no discernible impact on promoter activity *in vitro* (Supplementary Figure 5c) as may be expected considering the distance between Cbp1 binding site and promoter.

## Discussion

### Cbp1 functions as a facilitator of CRISPR array expression

Our results show that the coordinated action of two DNA-binding factors, the CRISPR array-specific Cbp1 and the general chromatin protein Cren7 facilitate CRISPR array function by specifically enhancing array transcription from the cognate leader promoters and suppressing transcription from cryptic promoters incorporated in CRISPR arrays (Figure 7). Thereby Cbp1 and Cren7 are preventing interference with CRISPR array expression e. g. by antisense transcription and overall safeguard the composition of the crRNA pool. Considering that the length of the leader-containing CRISPR arrays in *S. solfataricus* P2 (arrays A-E) appears to increase with enhanced Cbp1 and Cren7 binding, it is tempting to speculate that Cbp1 chromatinization allowed the emergence of the long CRISPR arrays characteristic for members of the order Sulfolobales genus that have Cbp1. Beyond a function of Cbp1 in array transcription, it may also protect the integrity of the genome, a function generally attributed to chromatin proteins, or more specifically improve the genetic stability of CRISPR arrays by suppressing recombination, which is a high risk due to the highly repetitive nature of CRISPR arrays. For example, Cbp1 binding to each repeat may inhibit homology search during homologous recombination events that could lead to the loss of spacers, and immunity. A recent study suggested that frequent recombination events in CRISPR arrays do occur, including in *S. solfataricus* P2 [21].

Cbp1 (and probably Cren7) binding to DNA does not interfere with all processes that utilise the DNA as a template. For instance, Cbp1 does not prevent spacer adaptation by the Cas1-Cas2 machinery *in vitro* [22]. Our occupancy mapping demonstrates that Cbp1 remains bound to CRISPR arrays during activation of CRISPR systems by SIRV2 infection in LAL14/1 (Figure 5), suggesting that the Cbp1-Cren7 chromatin is a constitutive part of the *Sulfolobales* CRISPR systems. But most importantly, Cbp1 binding enhances transcription of CRISPR arrays *in vivo* (Figure 3 and [7]) and in a minimal system *in vitro* (Figure 4), which may sound counterintuitive but shows that Cbp1-Cren7 does not form roadblocks for the RNAP and transcription elongation complex. Rather, Cbp1 is a positive factor that enables CRISPR expression, and it remains to be seen how this is integrated with other types of regulation. One previously identified regulator of CRISPR expression is the cyclic oligoadenylate-dependent transcription activator Csa3a. Csa3a is thought to control transcription directed by the leader promoters of CRISPR arrays C, D, and E in *S. solfataricus* P2 but not CRISPR arrays A and B, the two arrays with the highest Cbp1 chromatinization that have a deletion in the Csa3a binding site within the leader promoter [23].

### Abundant non-canonical Cbp1 binding sites in transposons

Our binding analyses identified many non-canonical, i. e. non-CRISPR, binding sites of Cbp1 in all three Sulfolobales strains (P2, REY15A, LAL14/1), including several strong binding sites in *S. solfataricus* P2 associated with intact or partial IS*C1229* transposons (Figure 1 and 6) that are not accompanied by Cren7 binding *in vivo*. Two distinct types of binding sites were present, one within the ORF of the transposase gene (n=2), and another one within the 3’-flank of the transposon (n=7). It is unlikely that these binding sites occur at random given that some of these binding sites show Cbp1 occupancy levels higher than CRISPR repeats with the exception of CRISPR A/B type repeats that are present in *S. solfataricus* P2 but not in *S. islandicus* strains [24]. IS*C1229* belongs to the IS*110* family of insertion sequences, which is characterized by the DEDD family transposase and a circular transposition intermediate formed through homologous recombination between the left and right flanks of the transposon. A curious feature of the IS*110* family transposons is that the strong promoter for transposase expression is formed only upon transposon circularization [25, 26]. If this mechanism is conserved in IS*C1229* transposons, Cbp1 binding in the 3’-flank of IS*C1229* could regulate either access of RNAP to the promoter (and hence transposase expression) or excision of the transposon. Notably, most of the IS*110* family elements in *S. solfataricus* and *S. islandicus* genomes are inactivated, decaying remnants, suggesting that proliferation of IS*110* is kept in check. In mammals, heterochromatin silences repetitive DNA elements and transposons, the role of Cbp1 binding in archaeal transposon function, including transposition remains to be investigated.

**Figure 6:**
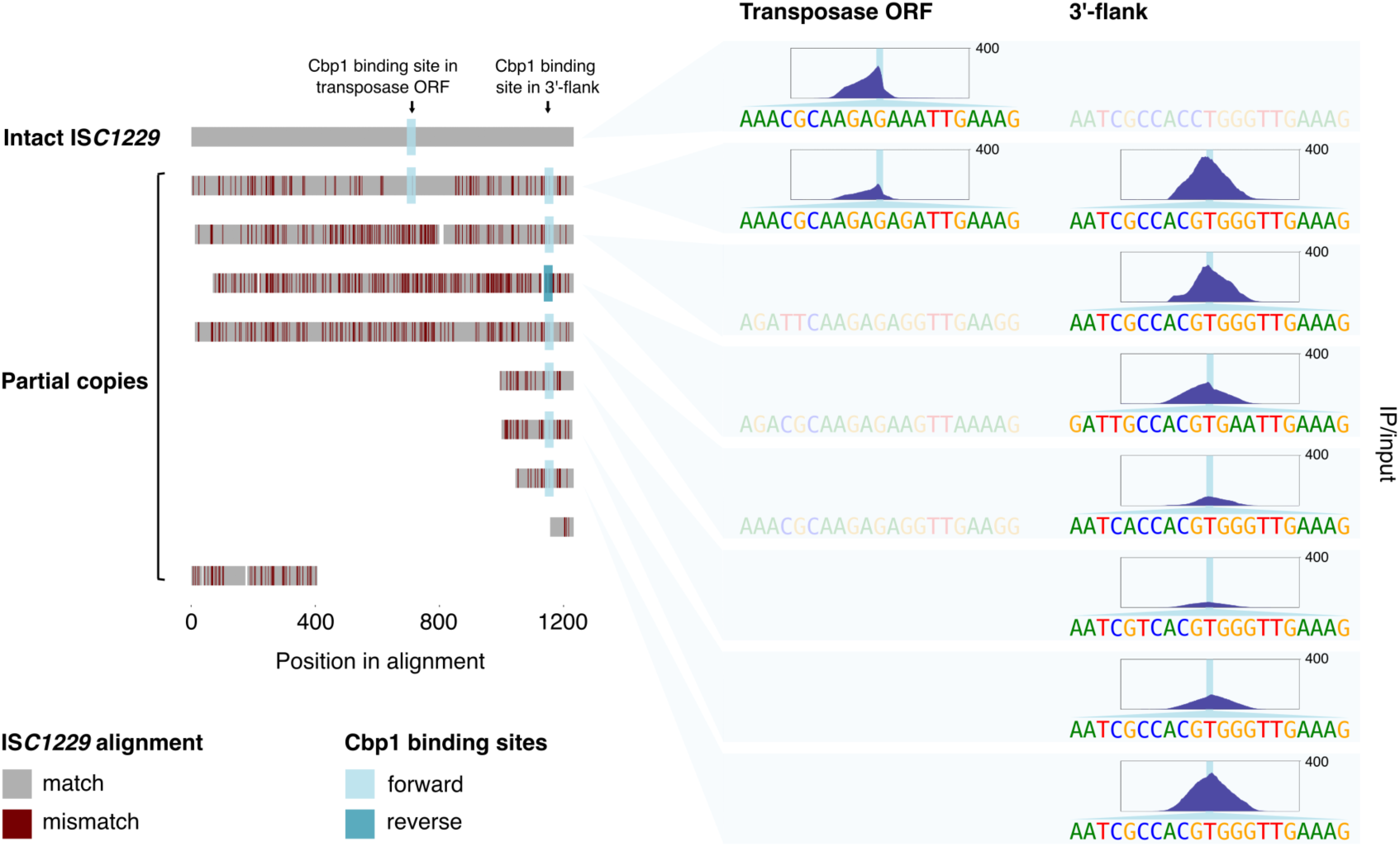
Cbp1 binds to IS110 family transposons. Alignment of IS*110* family transposons IS*C1229* in the *S. solfataricus* P2 genome. Partial IS*C1229* copies were aligned to the single full-length reference sequence with mismatches highlighted in dark red, insertions present in single IS*C1229* copies not shown. Cbp1 binding sites are highlighted by blue squares according to their orientation (defined by the orientation of Cbp1 binding on CRISPR arrays). The Cbp1 ChIP-seq occupancy at these binding sites is shown on the right (mean signal of two biological replicates) for a 500 bp centred around the 21 bp Cbp1 binding motif identified at each site. The sequence of the binding sites is shown below each ChIP-seq peak. The corresponding sequences in IS*C1229* copies that do not show Cbp1 binding are depicted as semi-transparent. Coordinates for intact and partial IS*C1229* copies were retrieved from [50]. Coordinates in the *S. solfataricus* P2 genome from top to bottom: 1745648-1746880 (intact copy serving as reference), 1449398-1450676, 1492743-1493952, 2818980-2820126,1761872-1763092, 1494821-1495060, 1486154-1486387, 2832377-2832564, 1761726-1761802, and 1476489-1476889.

### Evolution of chimeric Cbp1-Cren7 chromatinization

The non-canonical Cbp1 binding sites provided insights into the sequence dependence of Cbp1 binding, in particular they allowed us to identify a core motif of 9 bp which matches the 3’-end of CRISPR repeats (Figure 1b). This sequence is also conserved in CRISPR repeats found in Desulforococcales species which encode a Cbp1-related protein, Cbp2, that only consists of two HTH motifs [27]. Thus, the 9 bp motif likely represents the core binding site of both Cbp1 and Cbp2.

Cbp1 (and possibly Cbp2) coevolved with Cren7, a general chromatin protein with wider phylogenetic distribution and found in all Crenarchaeaota. Our data show that (i) Cbp1 recruits Cren7 *in vivo*, (ii) the Cbp1-Cren7-DNA complex features at least two Cren7 proteins recruited to the 3’ region of CRISPR repeats *in vitro*, and (iii) the third HTH domain of Cbp1 facilitates the Cren7 interaction (Figure 7). Multiple Cren7 interacting with the Cbp1-DNA complex or Cren7-Cren7 interactions could facilitate the binding of the second Cren7 molecule [28]. A cluster of multiple Cbp1 binding sites might be required to facilitate strong Cren7 recruitment *in vivo*, as the isolated, strong non-canonical Cbp1 binding sites are not associated with Cren7 enrichment in the genome, but the underlying differences in Cbp1 and Cren7 binding to CRISPR repeats and non-canonical binding sites remain to be solved. Heteromeric chromatin complexes are commonly formed by paralogous proteins such as in the octameric nucleosome complex in eukaryotes or the Alba-Alba2 complex in Crenarchaeota [29]. A heteromeric complex between two unrelated chromatin proteins in *Sulfolobales* that possess a large repertoire of chromatin proteins [5] points to a greater complexity of DNA chromatinization in archaea.

**Figure 7:**
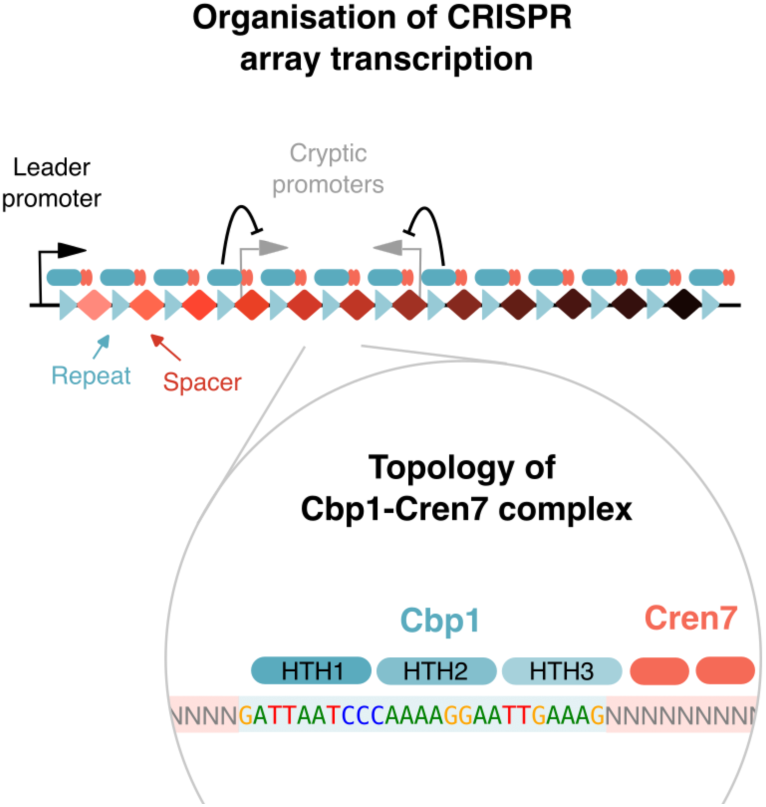
Model of Cbp1-Cren7 chromatinization and its role in CRISPR array transcription. Our data revealed the topology of the Cbp1-Cren7 complex. The three helix-turn-helix motifs of Cbp1 (HTH1 to 3) bind along the CRISPR repeat in 5’->3’ direction with HTH3 recruiting multiple Cren7 to the 3’-end of the repeat. Cbp1-Cren7 chromatinization alters CRISPR array transcription by blocking access of the transcription machinery to cryptic internal promoters while facilitating transcription from the leader promoter.

### The fluid boundaries between chromatin and transcription regulators

Several proteins cross the boundary between chromatin and transcription regulators, including H-NS in bacteria, TrmBL2 in *Thermococcus*, and Sa-Lrp and Lrs14 in Sulfolobales [5, 30]. Their regulatory mechanisms generally involve nucleation at high-affinity binding sites that is followed by spreading of the proteins along the DNA. While Cbp1 shows some unspecific DNA activity *in vitro* (see for example Supplementary Figure 4), our ChIP-seq data show that Cbp1 binding *in vivo* is limited to CRISPR repeats and other non-canonical Cbp1 binding sites. This remains true even for high affinity binding sites such as the IS*C1229* transposons and the CRISPR array 4 leader in *S. islandicus* LAL14/1. In this respect, chromatinization of CRISPR arrays is mediated by distinct high affinity sites similar to transcription regulators binding to operators but with the unique feature of multiple regularly spaced binding sites occurring in CRISPR arrays. The subsequent, or concomitant, recruitment of Cren7 is comparable to the spreading mechanism, expanding the chromatin structures beyond the CRISPR repeat.

The functioning of chromatin proteins, especially in eukaryotes, is commonly regulated by post-translational modifications, such as methylation and acetylation. In *S. islandicus*, chromatin proteins Alba1, Alba2, Cren7, Sul7d1 and Sul7d2 were found to be methylated [31], with the latter three proteins showing differential methylation patterns depending on the growth phase. Methylation on Lys50 has also been detected in Cbp1 [31]. It will be interesting to see whether post-translational modifications on Cbp1 and Cren7 affect their interaction with each other or with DNA.

In future, exploration of Cbp1-Cren7 chromatin in Sulfolobales will benefit from structural analyses of Cbp1-Cren7-DNA CRISPR arrays that will shed light on the topology of transcription unit-specific chromatin in archaea.

## Material and Methods

### Recombinant protein purification

The *S. solfataricus* P2 *cbp1* gene (SSO0454) was cloned into pET-21a(+) (Merck) vector and transformed into BL21 Star (DE3) cells (Thermo Fisher). Proteins were expressed in enriched growth medium for 3 hrs at 37°C after induction with 1 mM IPTG. Cells were resuspended in N200 buffer (25 mM Tris/HCl, pH 8.0, 10 mM MgCl_2_, 100 µM ZnSO_4_, 10% glycerol, 200 mM NaCl, 5 mM **β**-mercaptoethanol) and disrupted by sonication. Cell debris was removed by centrifugation. The lysate was incubated at 65°C for 10 min and centrifuged again. The heat-stable supernatant was loaded on a HiTrap Heparin column (GE Lifesciences) and eluted using gradient to 1 M NaCl. The peak fractions were combined, concentrated, and loaded onto a Superose 12 10/300 size exclusion column (GE Lifesciences) equilibrated in N200.

The *S. solfataricus cren7* gene (SSO6901) was cloned into pRSF-1b (Merck) and transformed into Rosetta2 (DE3) pLysS (Merck). 2 hrs after induction of expression with 1mM IPTG, cells were harvested and washed in 0.9% (w/v) NaCl. The protein was purified as described previously using a HiTrap cation exchange chromatography, heat incubation at 70°C for 30 min, heparin affinity and size exclusion chromatography in the final buffer (50 mM Tris/HCl (pH 8.0), 200 mM NaCl, 10 mM Na-EDTA, 10% glycerol, 10 mM **β**-mercaptoethanol) [32].

RNA polymerase and recombinant TBP, TFB were produced as described previously [33]. The recombinant core domain of TFB (TFB-C) was produced in the same manner as full-length TFB [14]. All protein concentrations were determined using the Qubit assay system (Thermo Fisher).

### Antisera and antibodies

Polyclonal rabbit antisera against recombinant *S. solfataricus* Cbp1 were produced at Davids Biotechnologie (Germany). Polyclonal rabbit antisera against *S. solfataricus* RNA polymerase (recombinant Rpo4/7 subcomplex), transcription factor TFB and sheep antiserum against Alba have been described previously [14, 33, 34]. The Alba antiserum was a kind gift of Malcolm White (University of St Andrews, UK). Protein G affinity-purified rabbit antibodies against recombinant *S. islandicus* Cren7 were purchased from CUSABIO (CSB-PA502491LA01FBP, Lot 00911A).

### Strains and *cbp1* gene deletion and cell growth

All genetic experiments were carried out in the *S. islandicus* REY15A strain E233S carrying deletions in the *pyrEF* genes conferring uracil auxotrophy as well as in the *lacS* gene [35]. ChIP-seq experiments were performed with a *cbp1* deletion strain described previously (strain 1) [35]. Since we were unable to revive the original *cbp1* deletion strain from glycerol stocks after prolonged storage time, we reconstructed *cbp1* deletion strains by a CRISPR-based strategy as previously described [36]. The protospacer adjacent motif (PAM) 5’-TCC-3’ (positioned at +150 referring to the start codon of *cbp*) and the 40-nt sequence immediately downstream of the PAM as protospacer were selected as targeting site. Spacer fragments were generated by annealing of the two complementary oligonucleotides KOcbp-SpF/ Kocbp-SpR and inserted into the artificial CRISPR array of pGE1 vector at the *Sap*I sites [37], yielding the interference plasmid pAC-*cbp1*. The donor DNA carrying mutant allele was generated by splicing with overlap extension PCR using the primers Kocbp-Lf/Kocbp-Lr and Kocbp-Rf/Kocbp-Rr. The resulting PCR products were inserted into the *Sph*I and *Xho*I restriction sites of pAC-*cbp1*, yielding the genome-editing plasmid pGE1-*cbp1*. The genome editing plasmid was then transformed into *S. islandicus* E233S competent cells by electroporation and transformants on the plates were validated by colony PCR with primers Kocbp-checkF/Kocbp-checkR. Curing of plasmid pGE1-*cbp1* was achieved by 5-Fluoroorotic acid counterselection. Two independent *S. islandicus* Δ*cbp1* strains (strains 2 and 3) were generated and verified by Sanger DNA sequencing. Strain 2 and 3 were used for TSS-RNA capable-seq experiments (see below).

All *S. islandicus* REY15A strains were grown in Brock medium [38] supplemented with 0.2% sucrose, 0.2% tryptone, and 20 µg/ml uracil at 75 °C in Erlenmeyer shake flasks at 150 rpm. *S. solfataricus* P2 was grown in Brock medium supplemented with 0.2% glucose, 0.1% tryptone likewise.

### SIRV2 propagation

An exponentially growing culture of *Sulfolobus islandicus* LAL14/1 [39] was infected with a preparation of *Sulfolobus islandicus* rod-shaped virus 2 (SIRV2). The infected culture was incubated at 76 °C under agitation for 2 days. After the removal of cells (7,000 rpm, 20 min; Sorvall 1500 rotor), viruses were collected and concentrated by ultracentrifugation (37,000 rpm, 2 h 30, 15 °C; Beckman Type 45 Ti fixed-angle rotor). The virus titer was determined using a plaque assay.

### Infection experiment and cross-linking

Four 250 mL cultures of *S. islandicus* LAL14/1 were grown in rich medium [19] at 76 °C under agitation for approximately 12 h. When the optical density (OD) reached 0.2, two of the cultures were infected with SIRV2 using a multiplicity of infection (MOI) of 10, while the other two served as uninfected controls. After 2 hours post-infection (hpi), 200 mL aliquots were rapidly transferred to flasks, placed on heated magnetic stirring plates and cross-linked with stabilized formaldehyde solution to a final concentration of 0.4%. The cross-linking proceeded for exactly 1 min and the reaction was stopped by adding Tris/HCl pH 8.0 to a final concentration of 100 mM. Samples were cooled down on ice for 5 min and centrifuged (7,000 rpm, 20 min, 4°C, Sorvall 1500 rotor). Cells were resuspended in 10 mL of PBS and pelleted using Eppendorf Centrifuge 5430 R (7,000 rpm, 20 min, 4°C). Pelleted cells were frozen in liquid nitrogen and stored at −80°C. Cell density and virus titer were measured at different time points to assess the efficiency of the infection.

### ChIP-seq and ChIP-exo experiments

ChIP-seq data for *S. solfataricus* RNA polymerase subunits Rpo4/7 and transcription factors TFB, Spt4/5, and aCPSF1 were obtained from NCBI GEO superseries GSE141290 [14]. All other ChIP-seq experiments were carried out using the same, previously described protocol [14, 40] with minor modifications as follows. *S. solfataricus* Cbp1, Rpo4/7, and TFB antibodies were purified from antiserum by Protein A agarose. All *S. solfataricus* antibodies and the *S. islandicus* Cren7 antibody showed good cross-reactivity in immunoprecipitation experiments for the other species. For *S. solfataricus* TFB and *S. islandicus* REY15A Cbp1, TFB, and Rpo4/7 ChIP experiments, 2 µg antibody were incubated with 500 µl cell lysate at 20 ng/µl DNA content. For Cren7 ChIP, the amount of antibody was increased to 4 µg. For ChIP experiments in *S. islandicus* LAL14/1, 2 µg Rpo4/7 or Cbp1 antibodies were incubated with 600 µl cell lysate at 12 ng/µl DNA content.

For ChIP-exo experiments, DNA shearing by sonication was altered to yield fragments within the recommended 200 – 1200 bp range. 1 ml of cell lysate was incubated with 8 µg Cbp1 or Cren7 antibody overnight at 4°C and antibodies were captured by further incubation with 50 µl Protein G-beads for 1 hr. Beads were washed six times in 1 ml RIPA buffer (50 mM HEPES/NaOH pH 7.6, 1 mM Na-EDTA, 0.7 % Na-Deoxycholate, 1% NP-40, 0.5 M LiCl) before two wash steps in 10 mM Tris/HCl pH 8.0. Library preparation was carried out following the ChIP-exo 5.0 method [41] with both the ExA1 and ExA2 adaptor sequences modified by introducing barcodes for dual indexing (ExA1_i5X_58: 5’-AATGATACGGCGACCACCGAGATCTACACN_8_ACACTCTTTCCCTACACGA CGCTCTTCCGATCT-3’ and ExA2_i7X: 5’-[Phos]-CAAGCAGAAGACGGCATACGAGATN_8_GTGACTGGAGTTCAGACGTGTGC TCTTCCGATCT-3’, where X and N denote the barcode number and sequence, respectively). Two biological replicates were used for all ChIP-seq and ChIP-exo experiments and deep-sequencing was carried out using Illumina HiSeq 125 cycle Paired-End Sequencing v4 or Hi-Seq 4000 75 Paired-End Sequencing.

### ChIP-seq data analysis

Mapping and normalisation of ChIP-seq data was conducted as previously described [14]. In brief, ChIP-seq paired-end reads were mapped using Bowtie v1.1.2 [42] with parameters -v 2 -m 1 –fr over the first 50 nt allowing only for uniquely mapped read pairs to be included. The mapped fragments were sampled to match a normal distribution with a mean of 120 and standard deviation of 18 or in the case of LAL14/1 ChIP-seq data a mean of 150, standard deviation of 20. Normalisation to input was carried out using deepTools bigwigCompare using signal extraction scaling (10,000 bins, 200 bp bin width [43].

### Cbp1 ChIP-seq peak calling

ChIP-seq peak calling was performed identically for each of the four different strains (*S. solfataricus* P2, *S. islandicus* REY15A, *S. islandicus* LAL14/1, and *S. islandicus* LAL14/1 + SIRV2) The paired-end alignment files for Cbp1 ChIP-seq data (see above) were converted to bed file format. Calling of narrow peaks on ChIP-Seq data was performed for each replicate using MACS2 version 2.2.6 [44] in the BEDPE mode with a cut-off of q = 0.01 and the call-summits subfunction activated. Input data were used as control sample. MACS2 estimated duplicate reads to be within the expected level. Peaks with summits located within the CRISPR arrays were removed for subsequent analysis. Matching of peaks between the two replicates was performed based on the peak summit positions differing by maximal 40 bp using BEDTools window [45]. *For S. solfataricus* P2 Cbp1, 199 matched peaks were further filtered for consistent ranking based on the p-value using the IDR method [46] with an estimated correlation coefficient rho = 0.99 for the reproducible component (global IDR < 0.01) representing a fraction of 0.80 of all peaks after 100 iterations. This resulted in a set of 120 peaks with a global IDR < 0.01. 92 peaks with an average enrichment of > 5 (based on MACS2 output) were used for all further data analysis (Supplemental file 1).

For *S. islandicus* REY15A Cbp1, IDR filtering resulted in an estimated correlation coefficient rho = 0.99 for the reproducible component (global IDR < 0.01) representing a fraction of 0.71 of all peaks after 100 iterations. This resulted in a set of 352 peaks with a global IDR < 0.01. 157 peaks with an average enrichment of > 5 were used for all further data analysis (Supplemental file 2).

For *S. islandicus* LAL14/1 Cbp1, IDR filtering resulted in an estimated correlation coefficient rho = 0.93 for the reproducible component (global IDR < 0.01) representing a fraction of 0.65 of all peaks after 100 iterations. This resulted in a set of 156 peaks with a global IDR < 0.01. 123 peaks with an average enrichment of > 5 were used for all further data analysis (Supplemental file 3).

For the SIRV2 infected LAL14/1 cells a lower enrichment threshold had to be applied because of the increased genomic DNA background in the ChIP saples. IDR filtering resulted in an estimated correlation coefficient rho = 0.91 for the reproducible component (global IDR < 0.01) representing a fraction of 0.52 of all peaks after 100 iterations. This resulted in a set of 94 peaks with a global IDR < 0.01. 86 peaks with an average enrichment of > 3 were used for all further data analysis (Supplemental file 4).

### DNA sequence motif identification

To identify DNA sequence motifs within the non-canonical binding sites of Cbp1, DNA sequences covering genomic intervals covering 80 bp (*S. solfataricus* P2) or 50 bp (*S. islandicus* REY15A) on either side of Cbp1 ChIP-seq peak summits (for peaks with minimum 5-fold enrichment) were retrieved using BEDTools getfasta [45]. Motif search was conducted using the MEME software version 4.11.2 with one occurrence of the motif per sequence [36]. A 0-order background model based on the respective genome sequences was employed.

### Cbp1 ChIP-exo data analysis

ChIP-exo reads were mapped using Bowtie v1.1.2 [42] with parameters -v 2 - m 1 –fr allowing only for uniquely mapped read pairs to be included. Bam files with alignments were filtered for first read in proper pairs (samtools view -f 66 -b) and the read 5’-end coverage per strand was calculated using BEDTools genomecov [45]. The geometric mean of 5’-end coverage for two biological replicates was calculated and scaled to 1x genome coverage. Aggregate plots were calculated for 198 CRISPR repeat in CRISPR arrays A and B and the 92 non-canoncial binding sites associated with Cbp1 ChIP-seq peaks.

### Cappable-seq

Cappable-seq data for *S. solfataricus* P2 were described in Blombach *et al.* 2021 [14] (NCBI GEO superseries GSE141290) and data for *S. islandicus* REY15A E233S (two biological replicates) and Δ*cbp1* strains 2 and 3 were generated likewise. In brief, cells were grown to exponential phase, cultures were mixed with 2 volumes of pre-cooled RNAprotect Bacteria Reagent (Qiagen), and cells were collected by centrifugation (5 min at 4000 × *g* at 4 °C). Small RNA preparations (20-200 nt length) were carried out using the mirVana miRNA isolation kit (Ambion/Thermo Fisher) following the manufacturer’s protocol. The library preparation including an enrichment step for 5’-triphosphorylated RNAs by capping the RNAs with 3’-desthiobiotin-TEG-GTP (NEB) [15] and subsequent deep sequencing on a Illumina NextSeq 500 system with 75 bp read length were conducted at Vertis Biotechnologie (Germany) as previously described [14].

### Cappable-seq data analysis

Data processing of *S. islandicus* REY15A Cappable-seq data was carried out as described previously [14]. In brief, poly(A)-tails and 3’-adaptors in the reads were removed, reads were aligned to the *S. islandicus* REY15A genome and the resulting bam files were merged and 5’-end coverage was calculated using BEDTools genomecov [45]. Bedgraph output was converted into bigwig file format.

For the identification of TSSs we proceeded as follows. We considered positions with an absolute increase of >20 read 5’-ends and a relative increase of >5-fold compared to the preceding position as candidate TSSs for each individual Cappable-seq sample and strand. Candidate TSSs reproducible across at least two samples (within or across conditions) were called. TSSs within the *cbp1* gene and its promoter were removed. To take into account more dispersed transcription initiation over a short region, we clustered called TSSs occurring within less than 5 bp distance and took the centre position of these TSS clusters as final TSS position. For the differential expression analysis, the Cappable-seq signal was integrated over a 11 bp window around the centre position of each TSS cluster and differentially expressed TSS s were called by Deseq2 [47] with a padj 0.01 cut-off.

### Differential binding analysis of *S. islandicus* REY15A Cren7

In order test for differential binding of Cren7 in *S. islandicus* REY15A E233S and the *cbp1* deletion strain1 ChIP-seq data, the CSAW package v1.26.0 [48] in R v.4.1.1 [49] was used for consecutive 30 bp windows and 250 bp maximal fragment length with a minimal count of 20 reads across the four libraries with two biological replicates per strain. A Benjamini-Hochberg multiple testing correction was applied for the differential binding results of the 30 bp windows. Windows overlapping with CRISPR arrays and non-canonical Cbp1 binding sites(peak summits from MACS2 peak calling) were obtained using BEDTools window [45] with maximal 30 bp distance (bedtool window -c -w 30).

### EMSA for templates with single Cbp1 binding sites

Double-stranded DNA templates bearing a 24 or 25 bp CRISPR repeat plus 20 bp flanking sequence on either side (or as otherwise indicated) were assembled by hybridisation of complementary oligonucleotides with one oligonucleotide being radiolabelled with [**γ**-^32^P]-ATP by T4 polynucleotide kinase (NEB). 15 µl EMSA samples contained 800 pM DNA template, 10 mM MOPS pH 6.5, 100 mM KCl, 10 mM MgCl_2_, 2 µg bovine serum albumin (NEB), 10% glycerol (v/v) and 5 mM DTT as well as in total 1.5 µl of Cbp1 and Cren7 preparations in N200 buffer. Samples were incubated for 5 min at 75°C and resolved on 6% native Tris-Glycine gels containing 2.5% glycerol and 1 mM DTT. Gels were dried and the signal was detected on BAS storage phosphor screens scanned on a Typhoon FLA 9500 scanner (GE Lifesciences).

### DNase footprinting assays

DNase I footprinting assay conditions were identical to those in EMSA experiments with the following modifications: the dsDNA concentration was raised to 10 nM with Cbp1 concentrations raised to 12.5 or 25 nM and Cren7 concentrations raised to 50 nM as indicated. After incubation for 5 min at 75°C, 15 µl samples were allowed to cool to room temperature and 1 µl RQ1 DNase (Promega) diluted to 0.1 U/µl in storage buffer (10 mM Hepes pH 7.5, 50% glycerol (v/v), 10 mM CaCl_2_, 10 mM MgCl_2_) was added to the samples. Samples were incubated for 10 min at room temperature before the addition of 16 µl formamide sample buffer (95% deionised formamide, 18 mM EDTA, 0.025% SDS). Samples were heated for 5 min at 95 °C before loading onto an 12% polyacrylamide, 7 M Urea, 1× TBE sequencing gel. Gels were dried and the signal was detected on BAS storage phosphor screens scanned on a Typhoon FLA 9500 scanner (GE Lifesciences).

### EMSA for templates with multiple Cbp1 binding sites

To test Cbp1 binding to the 600 bp DNA templates used in reconstituted *in vitro* transcription assays, we used identical buffer conditions and protein concentrations as in the transcription assays omitting nucleotides, RNA polymerase and transcription factors. Samples were incubated for 5 min at 65°C and 12 µl were loaded onto a 1.5% agarose gel. Gels were run for 16 hrs at ~0.9 V/cm in 1xTAE buffer and post-stained with ethidium bromide. Gels were visualised on a Typhoon FLA 9500 scanner (GE Lifesciences).

### Cross-linking of Cbp1 and Cren7

Cbp1 and Cren7 aliquots were buffer-exchanged into 25 mM HEPES/NaOH pH 7.5, 200 mM NaCl by ultrafiltration. DNA Templates were annealed in 10 mM HEPES/NaOH pH 7.5, 50 mM NaCl. Cbp1, Cren7, and template DNA concentrations were kept at 2.5 μM. Samples were incubated at 75°C for 5 minutes. 1 mM bissulfosuccinimidyl suberate (BS^3^) cross-linker (ThermoFisher) was added to each sample and incubated at 25°C for 30 minutes. Crosslinking reactions were quenched by the addition of 50 mM Tris/HCl (pH 8.0) and samples were resolved by SDS-PAGE followed by Coomassie Blue staining.

### Cell-free transcription assays

Templates for cell-free transcription were PCR-amplified as outlined in Supplemental file 5. *S. solfataricus* cell-free transcription assays were adapted from [14] for multi-round transcription conditions. 30 µl samples contained 20 µl cell lysate in 10 mM MOPS pH 6.5, 10 mM MgCl_2_, 2 mM spermidine supplemented with 10 mM rNTPs, trace amounts of [α-^32^P]-UTP and the indicated amounts of Cbp1 and Cren7. Samples were incubated at 70°C for 4 min before being placed on ice. Transcripts were isolated by affinity purification using 3’-biotinylated antisense oligonucleotides matching the first 25 nt of the transcripts (based on the mapped TSS, Supplemental file 5) as previously described [14] and resolved on 8% polyacrylamide, 7 M Urea, 1× TBE sequencing gels. Gels were dried and the signal was detected on BAS storage phosphor screens scanned on a Typhoon FLA 9500 scanner (GE Lifesciences).

### Reconstituted *in vitro* transcription assays

As template for reconstituted *in vitro* transcription assays, the initial 511 bp of *S. solfataricus* P2 CRISPR arrays A and B were amplified by PCR from genomic DNA (Supplemental file 5). The forward primer for the PCR encompassed a 46 nt sequence matching the strong viral model promoter T6 (positions −46 to −1 relative to TSS) fused to a priming region matching the initial 30 nt of the transcribed region of CRISPR array B. To test transcription through an inverted CRISPR array B (antisense direction), a corresponding construct was designed encompassing region −46 to +2 of the T6 promoter fused to the inverted 511 bp region of CRISPR array B. PCR products were cloned into vector pGEM-T (Promega). CRISPR A and B start naturally with a 6 nt cassette consisting of adenines and guanines that we used for synchronisation of transcription (see below). For the inverted CRISPR B construct, we altered the initial 6 nt to the same sequence using site-directed mutagenesis. All sequences were amplified by PCR from the plasmids using primers carrying 5 guanine nucleotides at their 5’-ends. Randomisation of the first repeat sequence in the T6 CRISPR B fusion was achieved by site-directed mutagenesis.

45 µl Transcription reactions containing 90 ng DNA template, 1 µg RNA polymerase, 1 µM TBP, 0.125 µM TFB, and the indicated concentrations of Cbp1 and Cren7 in transcription buffer (10 mM MOPS pH 6.5, 10 mM MgCl_2_, 105 mM KCl, 10% glycerol (v/v), 0.8 µg BSA, 10 mM DTT, 5 µg/ml heparin, 2mM spermidine) supplemented with 100 µM ATP/GTP were incubated for 5 min at 65 °C to allow for the formation of initially transcribing complexes. A single round of transcription was initiated by the addition of 250 µM ATP/GTP/CTP, 25 µM UTP supplemented with trace amounts of [α-^32^P]-UTP and 5 µM TFB core-domain variant. 10 µl were withdrawn at the indicated time points and mixed with 200 µl stop solution (0.3 M Na-Acetate pH 5.2, 10 mM Na-EDTA, 0.5% SDS, 150 µg/ml GlycoBlue (ThermoFisher). After extraction with Acid-Phenol-Chloroform (ThermoFisher) and ethanol precipitation, the pellets were dissolved in 10 µl formamide sample buffer (95% deionised formamide, 18 mM EDTA, 0.025% SDS) and heated for 5 min at 95 °C before loading onto an 8% polyacrylamide, 7 M Urea, 1× TBE sequencing gel. Gels were dried and the signal was detected on BAS storage phosphor screens scanned on a Typhoon FLA 9500 scanner (GE Lifesciences)

### Immunodetection of Cren7 in Δ*cbp1* strains

*S. islandicus* REY15A Δ*cbp1* strains 2 and 3 (2 biological replaces per strain) and the parental strain E233S (4 biological replicates) were grown to mid-exponential growth phase (O.D._600_ = 0.13 to 0.18) in 50 ml cultures. Cells were harvested by centrifugation and stored at −80°C. The cell material was resuspended in 500 µl TK150 buffer (20 mM Tris/HCl pH 8.0, 150 mM KCl, 2.5 mM MgCl_2_, 100 µM ZnSO_4_) supplemented with 50 µg/ml Dnase I and disrupted by sonication. After the removal of cell debris, the protein content of the lysate was estimated by using the Qubit protein assay (Thermo Fisher) and the lysate concentration was adjusted with TK150 buffer to 0.5 mg/ml. Samples were resolved using SDS-PAGE on a 14% Tris-Tricine gel and blotted onto a nitrocellulose membrane (GE Lifesciences). Cren7 was detected using a 1/1000 dilution of rabbit anti *S. islandicus* cren7 antibody (CUSABIO, Lot O0911A) as primary antibody and donkey anti-rabbit IgG Dylight680 (Bethyl Laboratories) as secondary antibody. As loading control, we detected the general chromatin protein Alba with sheep *S. solfataricus* Alba antiserum as primary antibody with donkey anti-sheep IgG Alexa488 (Thermo Fisher) as secondary antibody. Blots were scanned on a Typhoon FLA 9500 scanner (GE Lifesciences) and overlays were produced in ImageQuant TL 1D v8.2.0 (GE Lifesciences).

### Identification and alignment of IS110 family transposases

Transposon data were obtained from [50] and the ISfinder database [51]. Transposon sequences were aligned using MUSCLE [52] with insertions specific to single copies of the intact transposon removed. Fasta alignment files were imported into R and visualized using the msavisr function in the seqvisr package v0.2.7 (https://doi.org/10.5281/zenodo.6583981).

## Data availability

All sequencing data files (ChIP-seq, ChIP-exo, and Cappable-seq) and the processed data were deposited at NCBI GEO under accession code GSE226026.

## Code availability

The analysis code is available on github (https://github.com/fblombach/Cbp1_public)

## Acknowledgements

We thank Malcolm White (University of St Andrews, UK), Remus Dame (Leiden University, NL), Michael Terns (University of Georgia, GA) and Roger Garrett (University of Copenhagen, DK) for valuable advice on the project. Research in the RNAP laboratory at UCL is funded by a Wellcome Investigator Award in Science ‘Mechanisms and Regulation of RNAP transcription’ to FW (WT 207446/Z/17/Z).

## Supplementary tables and figures

**Supplementary table 1:** Genes within the exta ~28 kb deletion in the original Δcbp1 strain.

| GeneID | Old locus_tag | New_locus_tag | start | end | strand | annotation | remarks |
| --- | --- | --- | --- | --- | --- | --- | --- |
| 12417532 | SiRe_0633 | SIRE_RS03230 | 607599 | 608006 | - | IS200/IS605 family transposase |  |
| 12417533 | SiRe_0634 | SIRE_RS03235 | 608153 | 608584 | - | universal stress protein |  |
| 12417534 | SiRe_0635 | SIRE_RS03240 | 608901 | 609839 | - | peptidase |  |
| 12417535 | SiRe_0636 | SIRE_RS03245 | 609784 | 610980 | - | MFS transporter |  |
| 12417536 | SiRe_0637 | SIRE_RS03250 | 611017 | 611850 | - | KaiC domain-containing protein |  |
| 12417537 | SiRe_0638 | SIRE_RS03255 | 612007 | 612843 | + | undecaprenyl-diphosphatase |  |
| 12417538 | SiRe_0639 | SIRE_RS03260 | 612864 | 613799 | + | protease |  |
| 12417539 | SiRe_0640 | SIRE_RS03265 | 614065 | 614694 | + | GMP synthase |  |
| 12417540 | SiRe_0641 | SIRE_RS03270 | 614922 | 615443 | - | TQO small subunit DoxA domain protein |  |
| 12417541 | SiRe_0642 | SIRE_RS03275 | 615450 | 615971 | - | TQO small subunit DoxD |  |
| 12417542 | SiRe_0643 | SIRE_RS03280 | 616768 | 617958 | + | amidohydrolase family protein |  |
| 31508441 | NA | SIRE_RS13815 | 617959 | 618547 | - | permease | pseudogene, frameshift |
| 31508442 | NA | SIRE_RS13820 | 618557 | 619146 | - | hypothetical protein | pseudogene, partial |
| 12417545 | SiRe_0646 | SIRE_RS03300 | 619182 | 619715 | - | hypothetical protein |  |
| 12417547 | SiRe_0648 | SIRE_RS03305 | 620185 | 620883 | + | hypothetical protein |  |
| 12417548 | SiRe_0649 | SIRE_RS03310 | 621000 | 621803 | + | DNA repair ATPase |  |
| 12417549 | SiRe_0651 | SIRE_RS03315 | 621825 | 623201 | - | amino acid permease |  |
| 12417550 | SiRe_0650 | SIRE_RS03320 | 623184 | 623876 | + | hypothetical protein |  |
| 12417551 | SiRe_0652 | SIRE_RS03325 | 623989 | 625971 | + | acetyl-CoA synthetase |  |
| 12417552 | SiRe_0653 | SIRE_RS03330 | 626028 | 626558 | + | hypothetical protein |  |
| 12417553 | SiRe_0654 | SIRE_RS03335 | 626611 | 626925 | - | ferredoxin |  |
| 12417554 | SiRe_0655 | SIRE_RS03340 | 627074 | 627526 | + | hypothetical protein |  |
| 12417555 | SiRe_0656 | SIRE_RS03345 | 627554 | 627994 | + | hemerythrin |  |
| 12417556 | SiRe_0657 | SIRE_RS03350 | 628042 | 629085 | + | hypothetical protein |  |
| 12417557 | SiRe_0658 | SIRE_RS03355 | 629082 | 629810 | + | hypothetical protein |  |
| 12417558 | SiRe_0659 | SIRE_RS03360 | 630114 | 630356 | + | transcriptional regulator |  |
| 12417559 | SiRe_0660 | SIRE_RS03365 | 630371 | 631528 | - | peptidase U32 |  |
| 12417560 | SiRe_0661 | SIRE_RS03370 | 631643 | 632446 | - | enoyl-CoA hydratase/isomerase family protein |  |
| 31508443 | SiRe_0663 | SIRE_RS13825 | 632456 | 632638 | - | ring oxydation complex/ phenylacetic acid degradation | pseudogene, partial |
| 12417562 | SiRe_0662 | SIRE_RS03375 | 632622 | 632945 | + | transcriptional regulator |  |
| 12417563 | SiRe_0664 | SIRE_RS03380 | 632983 | 633669 | + | class II glutamine amidotransferase |  |
| 24772322 | NA | SIRE_RS03385 | 633723 | 633990 | + | hypothetical protein |  |
| 12417565 | SiRe_0665 | SIRE_RS03390 | 633943 | 635148 | - | transposase |  |

**Supplementary figure 1:**
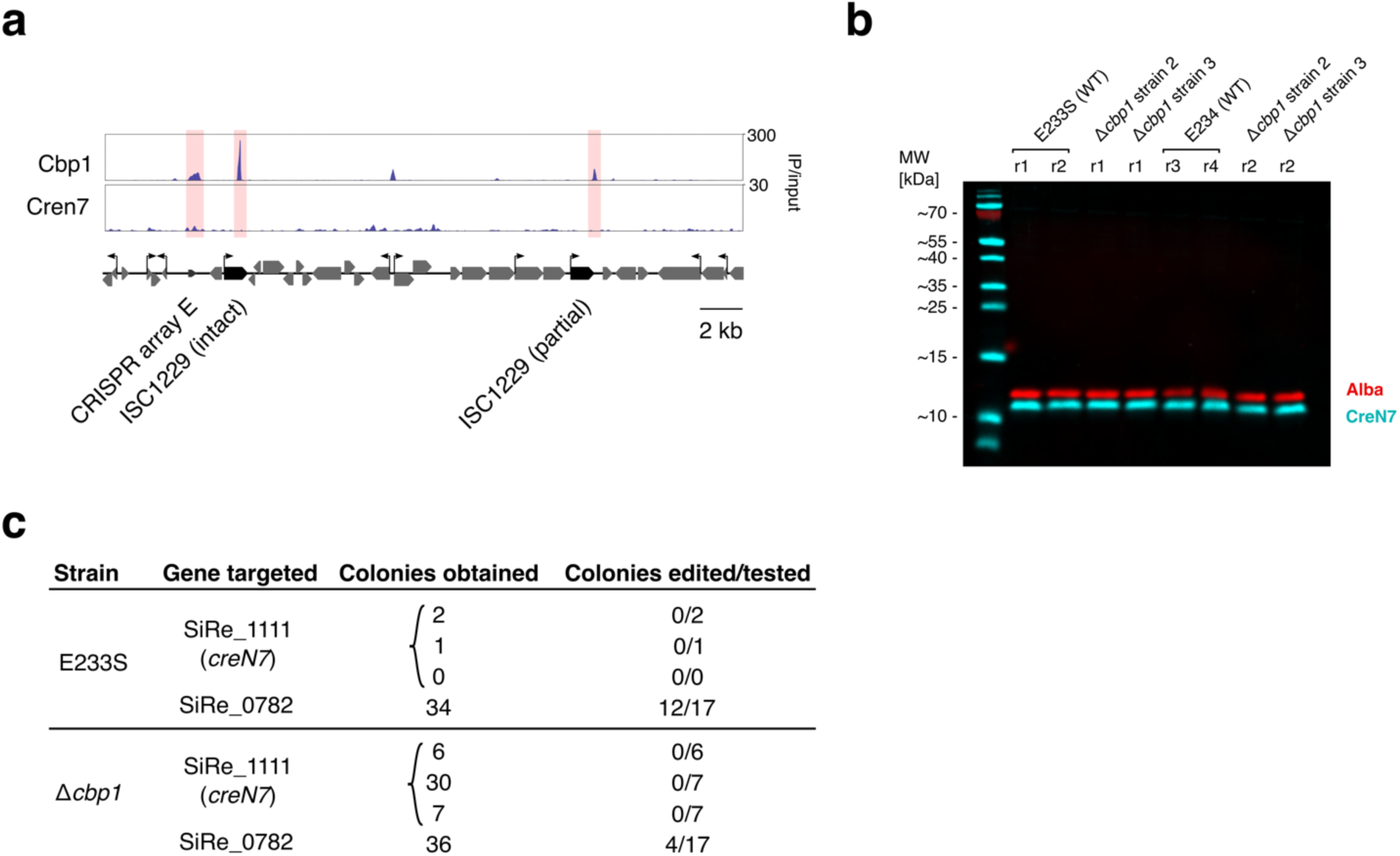
(a) Occupancy plot of Cbp1 and Cren7 in *S. solfataricus* P2 (ChIP-seq) showing discrete Cbp1 binding sites at ISC1229 transposons versus chromatinization at CRISPR array E. The mean ChIP-seq signal of two biological replicates is shown. (b) *cbp1* deletion does not affect Cren7 expression levels. Multiplex immunodetection of chromatin proteins Cren7 (cyan) and Alba (red, loading control) in parental strain E233S and two independent *cbp1* deletion strains (Δ*cbp1* strain 2 and strain 3). (c) Lethality of *cren7* loss does not depend on *cbp1.* Numbers for colonies obtained, tested and confirmed to be edited are shown for each independent transformation.

**Supplementary figure 2:**
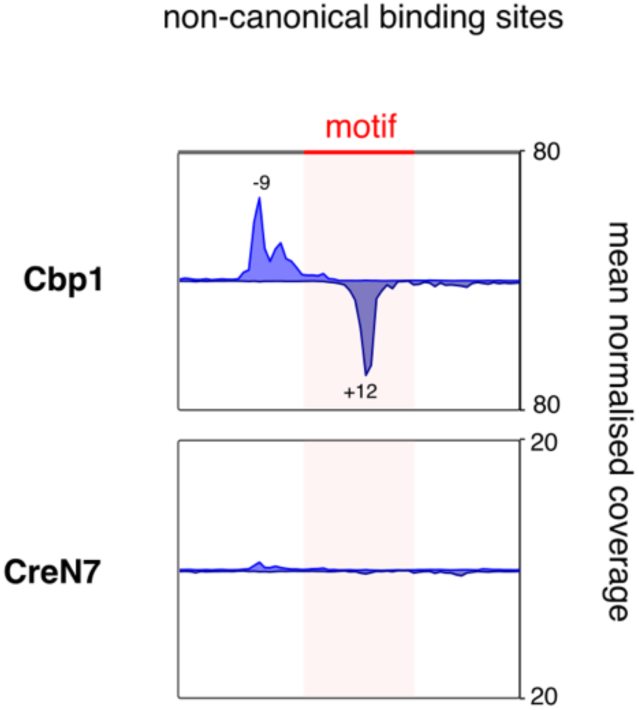
Cbp1 non-canonical binding sites are similar to CRISPR repeats in terms of ChIP-exo signal. Aggregate plots of Cbp1 and Cren7 ChIP-exo occupancy over 92 non-canonical binding sites in S. solfataricus P2. Signal for the plus and minus strand (relative to CRISPR array orientation) is plotted above and below the x-axis, respectively. The aggregate mean signal was calculated from the geometric mean of two biological replicates scaled to reads per million.

**Supplementary Figure 3.**
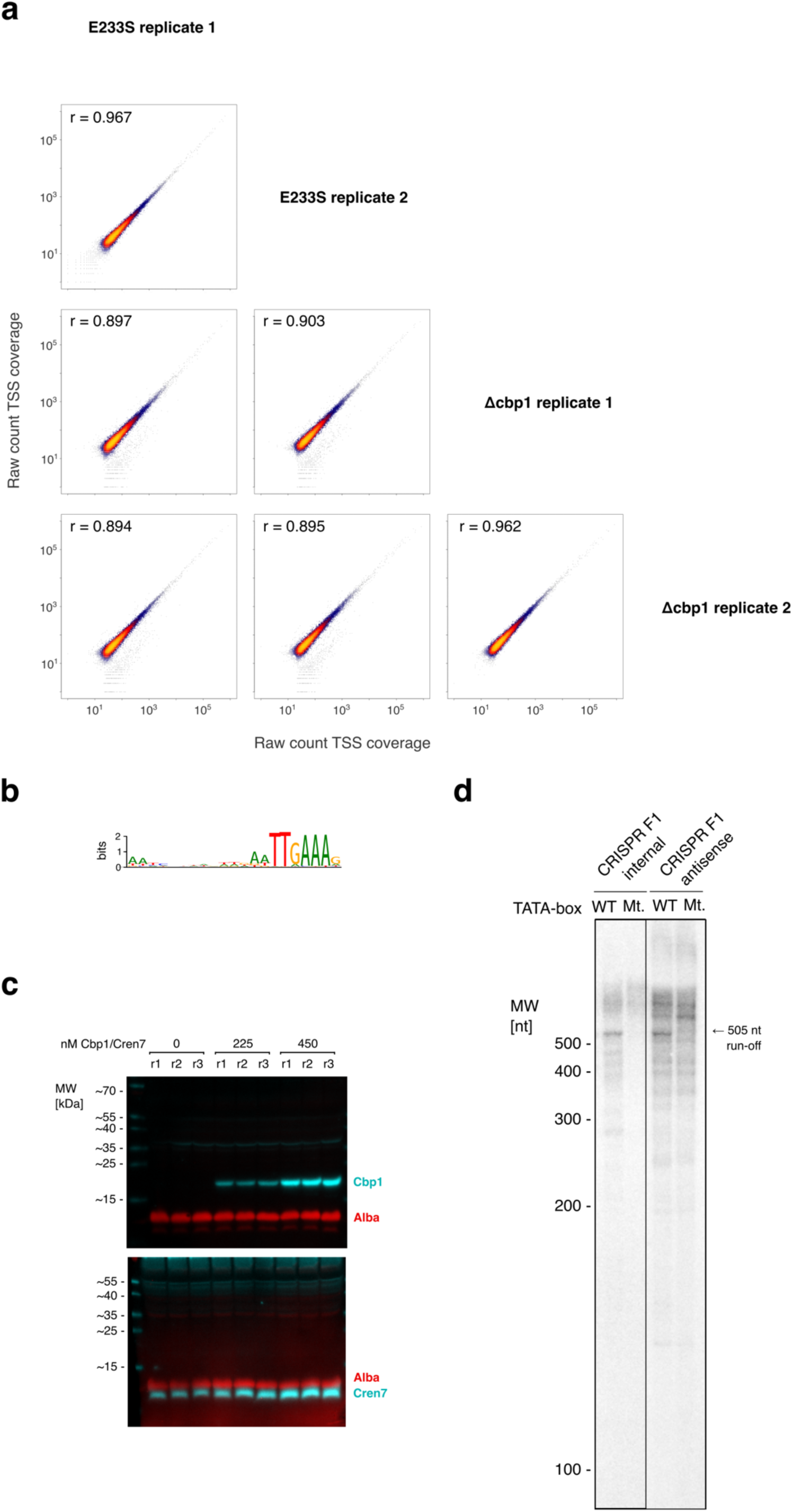
(a) Correlation of Cappable-seq replicates. The raw count data for 13150 called TSSs are plotted comparing the two E233S replicates and the two cbp1 deletion strains. The Spearman’s correlation coefficient between the samples is indicated on top. (b) MEME motif identified for 157 non-canonical binding sites of Cbp1 in *S. islandicus* REY15A. (c) Multiplex immunodetection of chromatin proteins Cbp1 and Cren7 (both cyan) and Alba (red, loading control) in *S. solfataricus* P2 cell lysate used for transcription assays. 225 nM or 450 nM recombinant Cbp1 and Cren7 were spiked into the cell lysate corresponding to 150 and 300 nM in the cell-free *in vitro* transcription reactions. Three technical replicates are shown. (d) Control cell-free transcription assay with TATA-box mutations in the internal and antisense promoters from CRISPR array F1.

**Supplementary Figure 4.**
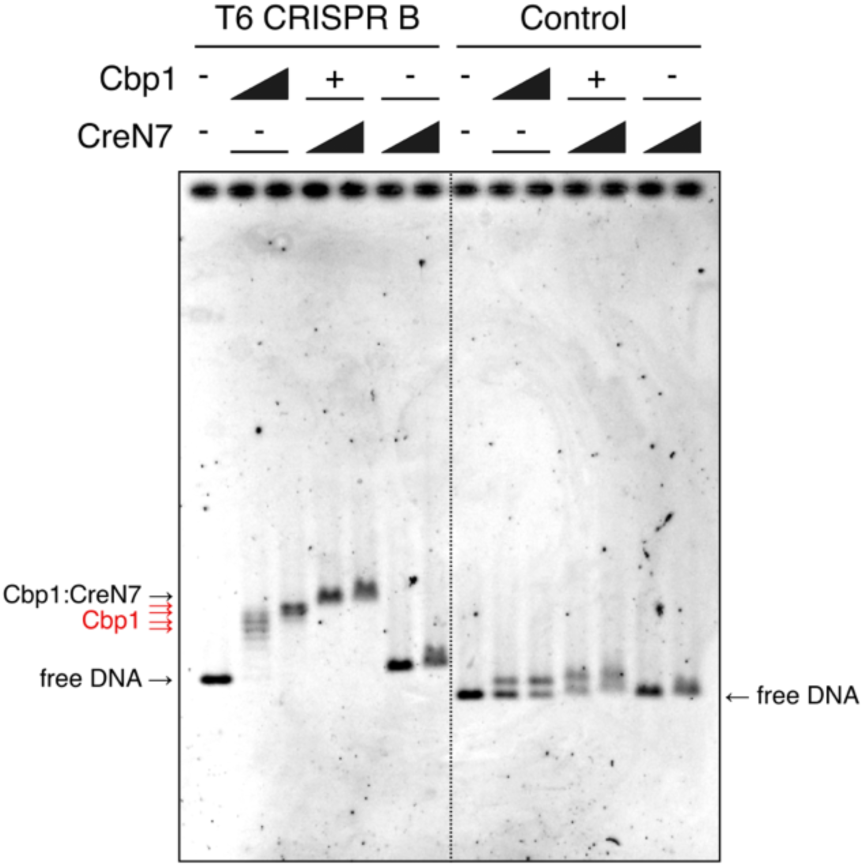
EMSA experiment testing Cbp1 and Cren7 binding to the T6 CRIPSR B template under in vitro transcription conditions (in absence of RNA polymerase, transcription factors and nucleotides). 150 or 300 nM Cbp1 and 300 or 600 nM Cren7 were tested in the reactions and resolved on a 1.5% agarose gel in 1x TAE buffer and post-stained with ethidium bromide. At lower Cbp1 concentrations, a ladder of bands reflecting recruitment of up to 8 Cbp1 to the CRISPR repeats was visible (highlighted in red) while binding saturates at 300 nM. A ~500 bp dsDNA derived from *Methanocaldococcus jannaschii* with similar AT-content served as control to test binding specificity of Cbp1.

**Supplementary figure 5.**
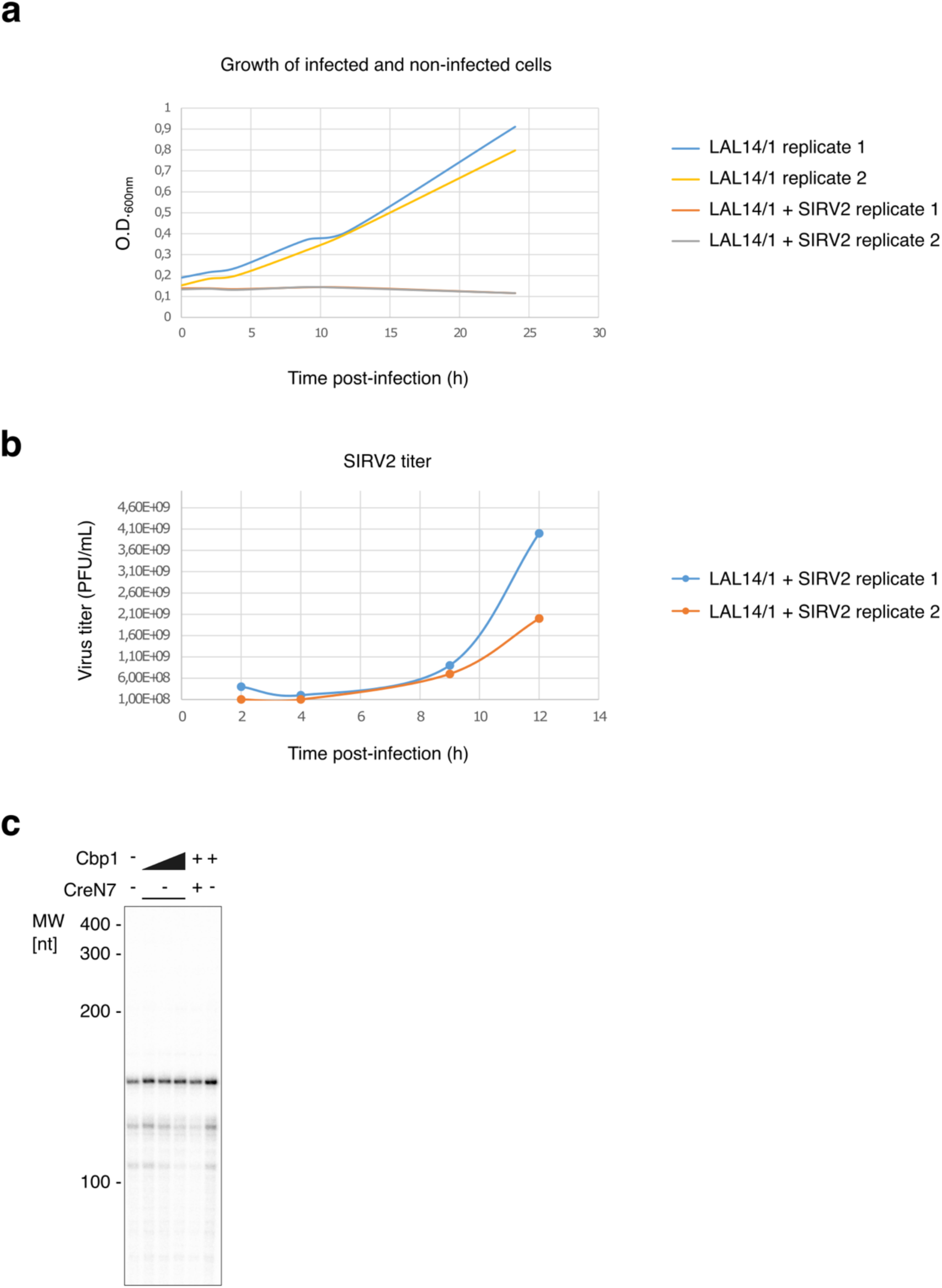
(a) Effect of SIRV2 infection on cell growth was monitored by measuring optical density at 600 nm for two biological replicates of uninfected and infected cultures, respectively. (b) SIRV2 titer over time after infection measured in PFU (plaque forming unit) per mL for the two infected cultures used in ChIP-seq experiments. (c) Reconstituted in vitro transcription assay on a template fusing the LAL14/1 CRISPR 4 promoter including the Cbp1 binding site with a synthetic C-less cassette under multi-round conditions. The nucleotide pool was restricted to ATP, GTP, and (radiolabelled) UTP to yield a ~150 nt transcript. 25, 100, 300 nM Cbp1 were tested alongside 300 nM Cren7 for their effect on transcription initiation.

